# Actinomycin D and bortezomib disrupt protein homeostasis in Wilms tumor

**DOI:** 10.1101/2024.06.11.598518

**Authors:** Patricia D.B. Tiburcio, Kenian Chen, Lin Xu, Kenneth S. Chen

**Author notes:** Correspondence: Kenneth S. Chen, UT Southwestern Medical Center, 5323 Harry Hines Blvd., Dallas, TX 75390.

## Abstract

Wilms tumor is the most common kidney cancer in children, and diffuse anaplastic Wilms tumor is the most chemoresistant histological subtype. Here, we explore how Wilms tumor cells evade the common chemotherapeutic drug actinomycin D, which inhibits ribosomal RNA biogenesis. Using ribosome profiling, protein arrays, and a genome-wide knockout screen, we describe how actinomycin D disrupts protein homeostasis and blocks cell cycle progression. We found that, when ribosomal capacity is limited by actinomycin D treatment, anaplastic Wilms tumor cells preferentially translate proteasome components and upregulate proteasome activity. Based on these findings, we tested whether the proteasome inhibitor bortezomib sensitizes cells to actinomycin D treatment. Indeed, we found that the combination induces apoptosis both *in vitro* and *in vivo* and prolongs survival in xenograft models. Lastly, we show that increased levels of proteasome components are associated with anaplastic histology and worse prognosis in Wilms tumor patients. In sum, maintaining protein homeostasis is critical for Wilms tumor proliferation, and it can be therapeutically disrupted by blocking protein synthesis or turnover.

## INTRODUCTION

Wilms tumors, or nephroblastomas, are the most common pediatric kidney cancer. Globally, Wilms tumor is diagnosed in 10.4 per 1 million children less than 15 years old each year ^1^. North American risk stratification criteria classify Wilms tumors as having favorable or anaplastic histology. The development of effective combinations of chemotherapy, radiation, and surgery has pushed the 5-year overall survival rate to over 90% for those with favorable histology Wilms tumor (FHWT). These strong cure rates have allowed us to de-intensify therapy for some patients with FHWT, where up to 24% of long-term survivors who were treated on historical regimens developed therapy-related chronic health conditions ^2^. Other FHWT patients receive intensified therapy based on biological risk factors such as combined loss-of-heterozygosity (LOH) of chromosomes 1p and 16q or gain of chromosome 1q. On the other hand, patients with diffuse anaplastic Wilms tumor (DAWT), which accounts for ∼10% of Wilms tumor patients, continue to have a relapse rate of over 40% and a 4-year overall survival rate of 66% despite being treated with more aggressive drugs such as doxorubicin, etoposide, cyclophosphamide, and carboplatin ^1,3–5^. Recent attempts to intensify chemotherapy for those with DAWT have increased short-term toxicity with minimal improvements in cure rate ^6^. Anaplastic histology is associated with loss or mutation of p53 ^7,8^, but the molecular mechanisms underpinning chemoresistance in anaplasia are unknown, and there remain no targeted therapies effective in Wilms tumor.

Since the 1960s, chemotherapy regimens utilizing actinomycin D (actD), also called dactinomycin, have been routinely used to treat Wilms tumors ^9,10^, rhabdomyosarcoma, and other sarcomas ^11,12^. In DAWT, however, actD has been removed from standard protocols. Unlike most FHWT regimens, DAWT patients receive doxorubicin, which can provide higher rates of cure but carries more short- and long-term toxicities, including cardiotoxicity and secondary cancers ^13^. Understanding how to improve actD sensitivity without the toxicities associated with anthracyclines like doxorubicin could improve outcomes for Wilms tumor as well as other malignancies.

The molecular activity of actD is concentration-dependent. At high concentrations, it is a DNA-intercalating agent that can block transcription and DNA replication. However, at the low-nanomolar serum concentrations typically achieved in patients, it primarily inhibits transcription of ribosomal RNA (rRNA), which is transcribed by RNA polymerase I (Pol I), and transfer RNA (tRNA), which is transcribed by RNA polymerase III (Pol III) ^14–18^. Ribosomes are composed of rRNA and ribosomal proteins (RP), and impaired Pol I activity results in fewer fully formed ribosomes, which consequently reduces global translation. In ribosome-depleted settings, the remaining ribosomes are not uniformly distributed across remaining transcripts; instead, they favor certain transcripts, based on factors such as their intracellular localization, secondary structure, and presence of specific sequence motifs ^18,19^ (Figure 1A). We thus reasoned that the genes preferentially translated following actD exposure could be therapeutically targetable in anaplastic Wilms tumor. To date, however, no published findings have characterized how actD affects translation in Wilms tumors.

**Figure 1.**
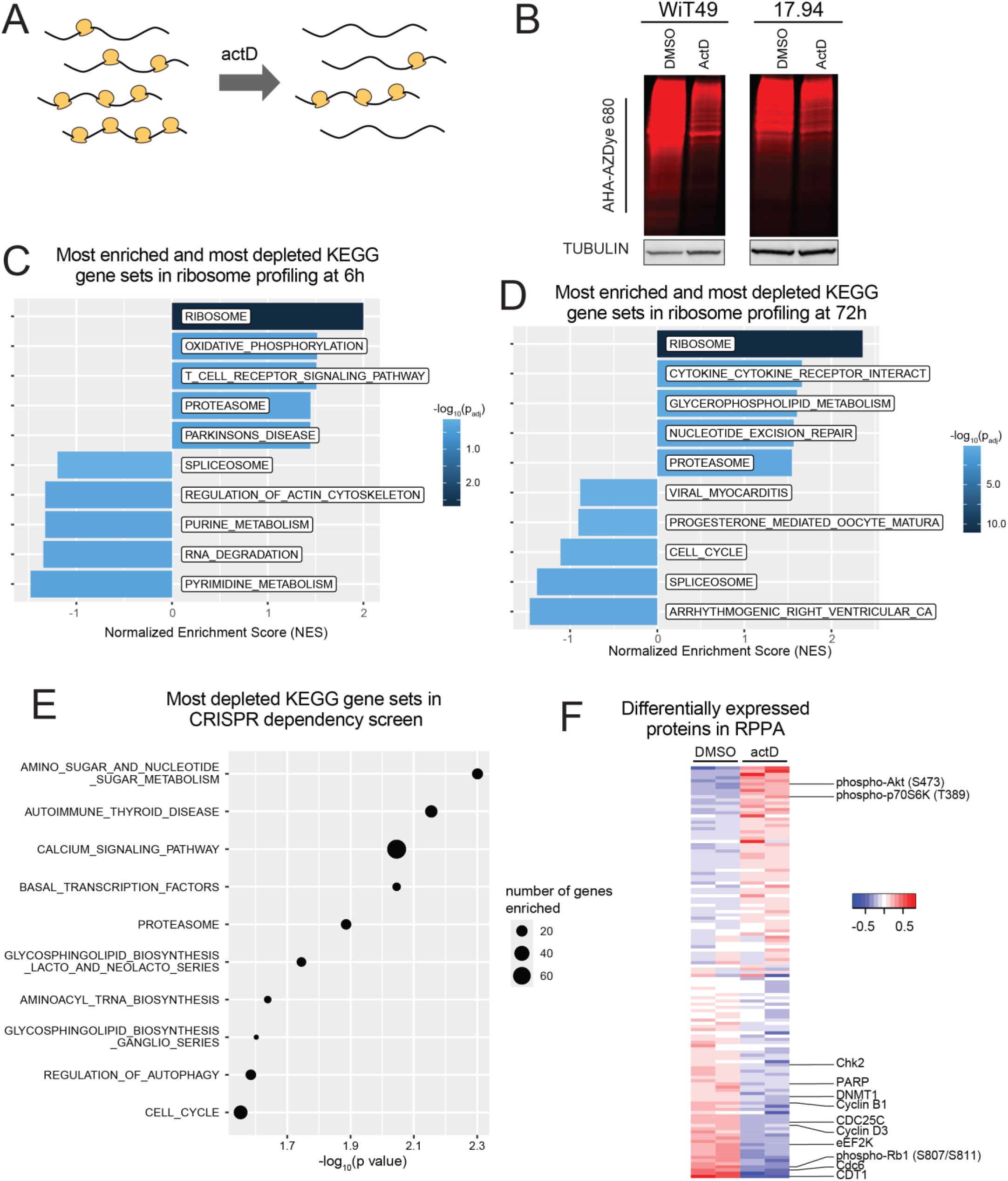
ActD disrupts protein homeostasis in anaplastic Wilms tumor cells. **(A)** Diagram of the effect of actD at the translational level. ActD depletes fully-formed ribosomes and reduces translational capacity. The reduced ribosomes are distributed unevenly. Ribosome profiling provides a snapshot of transcripts actively undergoing translation. **(B)** Whole protein lysates of DMSO (Vehicle) or actD-treated WiT49 or 17.94 fluorescently labeled with AHA-AZDye 680 for nascent protein detection and corresponding loading control TUBULIN detected by western blot from the same gel. **(C,D)** Top 5 most enriched and bottom 5 most depleted KEGG gene sets in ribosome profiling of actD- vs. DMSO-treated WiT49 after 6 hours **(C)** or 72 hours **(D)**. **(E)** GSEA of genome-wide CRISPR knockout screen reveals top 10 most significant KEGG pathways that sensitize WiT49 to actD, ranked by p-value. **(F)** Heatmap displaying RPPA results of actD- vs. DMSO-treated WiT49. Normalized, log2-transformed, median-centered values from validated antibodies with standard deviation over 0.1 are shown here.

In this study, we used ribosome profiling to find that proteasome components are preferentially translated in anaplastic Wilms tumor cell lines following actD treatment. Based on these findings, we studied a combination with the proteasome inhibitor bortezomib (BTZ), and we found that it increases sensitivity of anaplastic Wilms tumor cells to actD *in vitro* and *in vivo*. Lastly, DAWTs express higher transcript levels of several proteasome components than relapsed FHWTs, and higher levels of proteasome components are associated with worse prognosis.

## RESULTS

### Actinomycin D alters the translational landscape of anaplastic Wilms tumor cells

ActD blocks rRNA transcription at nanomolar doses and mRNA transcription at micromolar doses ^18^. To examine the effect of actD on cell viability at nanomolar doses in Wilms tumor, we measured 72-hour actD sensitivity in two anaplastic Wilms tumor cell lines, WiT49 and 17.94 (Supplementary Figure S1A). For these cell lines, we measured IC50 to be 1.3 and 2.2 nM, respectively. Next, we confirmed that 2 nM actD reduces levels of 45s pre-rRNA and 18s mature rRNA in both cell lines (Supplementary Figure S1B). This results in decreased overall protein synthesis, consistent with specific impairment of Pol I activity, as measured by incorporation of both the methoinine analog L-azidohomoalanine and O-propargyl-puromycin (Figure 1B, Supplementary Figure S1C, S1D).

Thus, to understand how actD affects protein levels and cellular functions in Wilms tumor, we next performed three complementary assays in parallel: ribosome profiling, to identify preferentially translated transcripts; reverse-phase protein arrays (RPPA), to identify differences in protein levels and post-translational modifications; and a CRISPR dropout screen, to identify targetable vulnerabilities.

First, to ascertain preferentially translated transcripts, we used ribosome profiling (also known as ribosome footprinting), which provides a snapshot of the transcripts actively undergoing translation ^20,21^. This technique entails trapping ribosomes on mRNA transcripts, degrading unprotected RNA, and sequencing the RNA fragments that were protected from degradation by ribosomes. Sequencing of the ribosome-protected RNA is computationally compared to RNA sequenced conventionally in the same samples to calculate “translational efficiency”, a measure of the active translation of each transcript.

Specifically, we performed ribosome profiling in WiT49 cells treated with actD or vehicle control at 6 hours, 72 hours, and two weeks. At each timepoint, we performed ribosome profiling to quantify gene-level differences in translational efficiency in actD-versus vehicle-treated cells. To confirm the expected effect of depleting ribosomes, we examined the locations of the ribosome footprint edges in metagene plots around translation start sites (Supplementary Figure S2A-S2F). As expected, in both DMSO- and actD-treated cells, there were essentially no detectable reads in 5’ untranslated regions (5’UTRs), while coding sequences exhibit a trinucleotide periodicity, reflecting the reading frame of elongating ribosomes. In DMSO-treated cells, the left edge of ribosome footprints accumulated just upstream of translation start sites, which reflects pausing of the ribosome at the initiation site, as expected for normal conditions (Supplementary Figure S2A-S2C). In actD-treated cells, however, the initiation site peak was blunted, suggesting that when ribosomes are depleted, the ribosomes that remain spend less time paused at initiation (Supplementary Figure S2D-S2F). For all durations of treatment, actD appeared to have minimal effect on the transcriptome, while ribosome footprints revealed gross perturbation of translational landscapes by actD (Supplementary Figure S3A-S3B). After two weeks of intermittent actD dosing, cells appear to return to a new steady state with globally reduced translation.

We next compared the translational efficiency of each gene in actD-treated cells at 6 and 72 hours, and we connected preferentially translated genes into Kyoto Encyclopedia of Genes and Genomes (KEGG) pathways using GSEA (Supplementary Table S1). The most enriched KEGG pathway by translational efficiency at both timepoints was KEGG_RIBOSOME, which is composed of RPs and genes that regulate ribosome biosynthesis (Figures 1B, 1C; Supplementary Figures S4A, S4B; Supplementary Table S2). This is consistent with reports that RPs are unstable when actD depletes rRNA, which can upregulate translation of RPs via mTORC1 signaling ^22,23^. This increase in translational efficiency was accompanied by a relative decrease in mRNA abundance for ribosomal proteins (Supplementary Figure S4C). KEGG_PROTEASOME was the only other gene set in the top five most preferentially translated gene sets at both timepoints (Figure 1C, 1D; Supplementary Figure S4D, S4E; Supplementary Table S2). This increase in translational efficiency was also accompanied by a relative decrease in mRNA abundance for proteasomal proteins (Supplementary Figure S4F). On the other hand, the most downregulated gene sets at 6 hours entailed nucleotide turnover, which led to a depletion of cell cycle genes at 72 hours (Figure 1C, 1D; Supplementary Table S2). In sum, our ribosome profiling of actD-treated cells detected preferential translation of RPs and proteasome components, with a concomitant decrease in translation of cell cycle genes.

Next, we used a genome-wide CRISPR screen to identify therapeutic vulnerabilities in cells with intermittent actD or DMSO for 14 days. We again used KEGG pathways to categorize dependency genes (Figure 1E; Supplementary Table S3). Here, we again found enrichment for pathways related to protein turnover, including KEGG_PROTEASOME, as well as nucleic acid turnover and cell cycle.

Thirdly, since actD regulates protein synthesis, we also performed RPPA on WiT49 cells treated with actD or DMSO for 72 hours to understand how actD affects protein levels and post-translational modifications (Figure 1F; Supplementary Table S4). We found an increase in phosphorylation of some components of the mTORC1 signaling pathway, which mediates the feedback signaling to upregulate translation of RPs in the setting of rRNA depletion ^23^. On the other hand, cell cycle markers such as CDT1, CDC6, and phosphorylated RB1 were depleted in actD-treated cells (also see Figure 3D below). This is consistent with depletion of cell cycle gene sets in ribosome profiling at 72 hours. Together, our three datasets show that, in the presence of actD, Wilms tumor cells choose to preferentially translate ribosome and proteasome components rather than progress through the cell cycle.

### mTORC1 inhibition does not sensitize Wilms tumor cells to actinomycin D

Based on phosphorylation of mTORC1 signaling intermediates in RPPA results and prior interest in mTORC1 signaling in Wilms tumor ^24,25^, we next investigated how actD affected mTORC1 signaling in WiT49 and a second anaplastic Wilms tumor cell line, 17.94. Using Western blots, we confirmed that actD induced phosphorylation of AKT and 4E-BP1 in WiT49 and 17.94 cells (Supplementary Figure S4G, S4H). (The phosphorylation of another mTORC1 target, p70S6K, was not clearly upregulated by actD.) We treated WiT49 and 17.94 with rapamycin at doses up to 100 µM and found that this drug confers a more cytostatic rather than cytotoxic effect (Supplemental Figure S4I). We also measured cell viability in combinations of actD and the mTORC1 inhibitor rapamycin using the Loewe independence model (Supplementary Figure S4J). However, the interaction did not consistently show synergy, as we only observed synergy above the upper limit of serum concentrations typically achieved in patients (∼2-3 nM for actD^14^ and 16 nM for rapamycin^26^). In other words, mTORC1 inhibition with rapamycin did not appear to consistently sensitize Wilms tumor cells to actD, and rapamycin alone was not cytotoxic, even at very high doses.

### ActD induces proteotoxic stress and cell cycle arrest in anaplastic Wilms tumor cells

One effect of mTORC1 signaling is to upregulate translation of RPs ^27^, and KEGG_RIBOSOME was the most enriched gene set in actD-treated cells. However, we found that 72-hour actD treatment in fact appeared to slightly reduce the total protein levels of multiple RPs in both WiT49 and 17.94 (Supplementary Figure S5A, S5B). This could be because excess RP subunits are unstable without rRNA and are degraded by the proteasome to maintain appropriate RP:rRNA stoichiometry ^22,28,29^.

While we did not observe an increase in RP levels with actD treatment, we did observe accumulation of proteasome components. When we exposed WiT49 and 17.94 to actD for up to 72 hours, both cell lines exhibited an appreciable increase in several proteasome complex subunits at one or more timepoints between 18 and 72 hours (Figure 2A, 2B). We specifically measured PSMD1 (also known as P112), PSMA6 (subunit α1), PSMB1 (β6), PSMB2 (β4), and PSMB5 (β5), as well as the molecular chaperone for proteasome assembly, POMP. In both cell lines, PSMA6, PSMD1, and POMP peaked at 18-24 hours. PSMB1 peaked at 72h in WiT49 and but rose earlier in 17.94 and remained high, while PSMB2 peaked at 24h in 17.94 but rose earlier in WiT49 and remained high. Lastly, PSMB5 peaked at 48-72 hours in WiT49 but did not appear to rise in 17.94 at the timepoints we examined. These increases in protein levels of proteasome components corresponded to increased proteasome chymotrypsin-like enzymatic activity in both cell lines, which was blunted by co-treatment with the proteasome inhibitor bortezomib (BTZ, Figure 2C).

**Figure 2.**
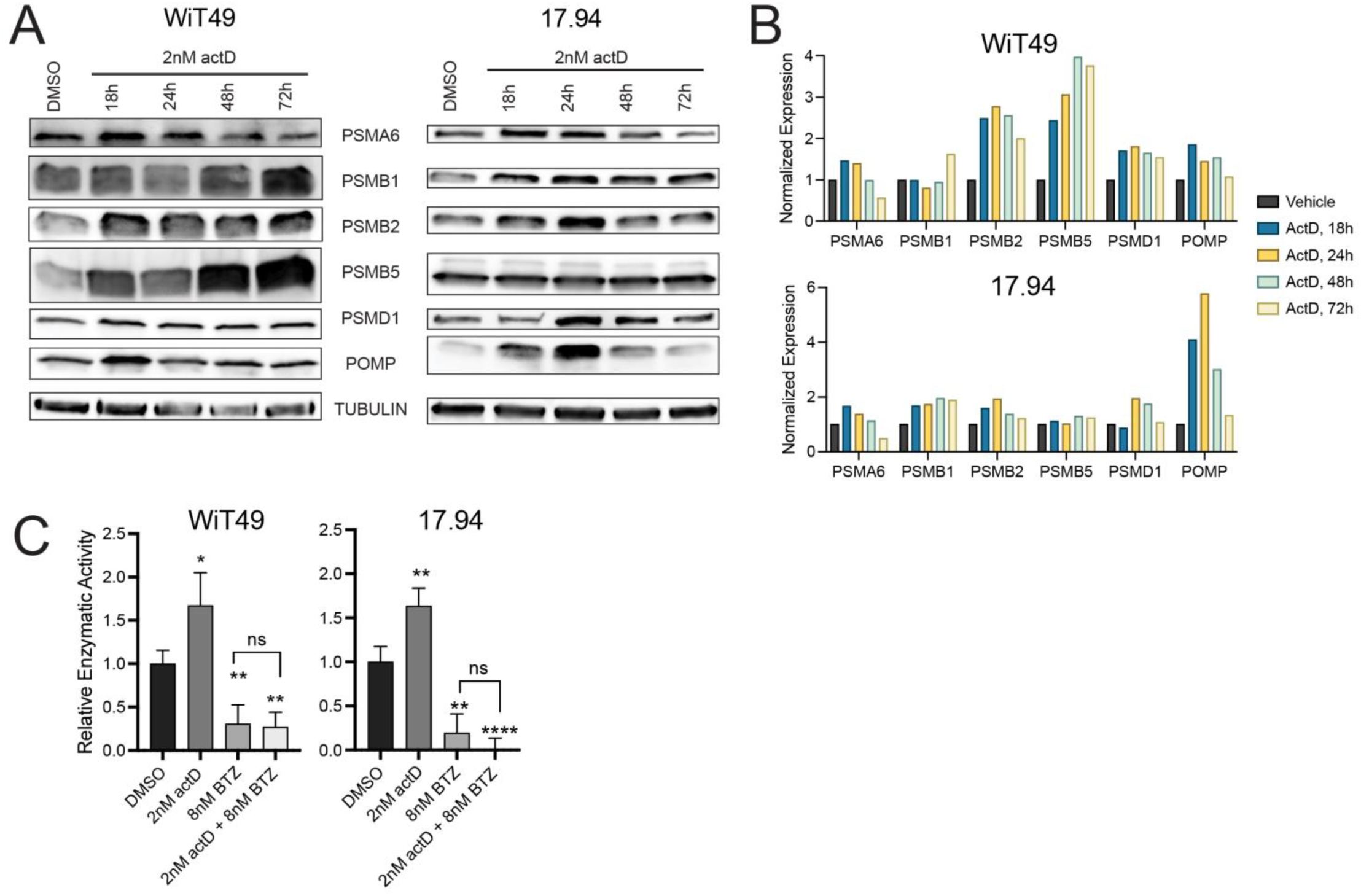
ActD promotes proteasome level and activity in anaplastic Wilms tumor cells. **(A,B)** Western blots for **(A)** and quantification of **(B)** proteasome subunits PSMA6 (α1), PSMB1 (β6), PSMB2 (β4), PSMB5 (β5), and PSMD1 (P112), and POMP in WiT49 and 17.94 following 18-, 24-, 48-, and 72-hour 2nM actD treatment versus vehicle (DMSO). **(C)** Quantification of relative proteasome-specific chymotrypsin-like activity in WiT49 and 17.94 cells following 24 hours of DMSO versus 2nM actD, 8nM BTZ, or 2nM actD + 8nM BTZ treatments (Student’s t-test p-value versus vehicle: *<0.05, ** <0.01, ****<0.0001).

Because actD caused preferential translation of proteasome subunits, we next examined how WiT49 and 17.94 respond to proteasome inhibition. These two cell lines were sensitive to BTZ alone at nanomolar concentrations, in the range of serum levels typically achieved in patients ^30,31^ (Figure 3A). This is close to the average sensitivity (3.9 nM) observed across the NCI-60 cell lines screen ^32^. Furthermore, at or below these levels, actD and BTZ synergistically inhibited WiT49 and 17.94 to different degrees (Figure 3B). In WiT49, actD and BTZ acted synergistically across nearly all combination concentrations. Loewe scores in 17.94 showed synergy at lower concentrations and additivity at medium and higher concentrations. Taken together, these data suggest that the addition of BTZ could be a strategy to enhance or restore actD sensitivity in anaplastic Wilms tumor cells.

**Figure 3.**
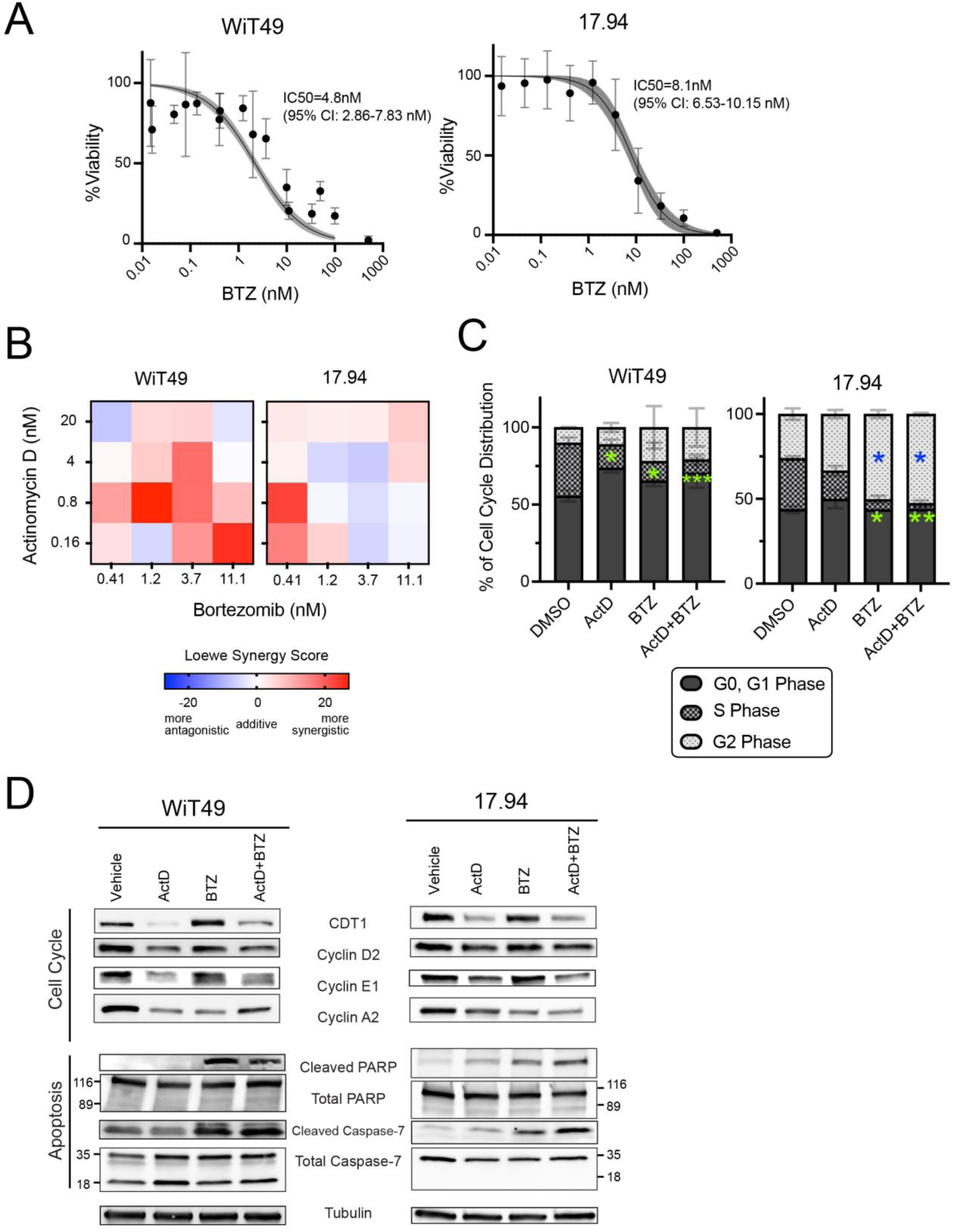
BTZ sensitizes anaplastic Wilms tumor cell lines to actD *in vitro*. **(A)** BTZ kill curves for WiT49 and 17.94 with IC50 values indicated (four-point regression line with shaded region representing 95% confidence interval). **(B)** Heat maps of Loewe synergy scores for combinations of actD and BTZ in WiT49 and 17.94 cells. **(C)** Distribution across phases of the cell cycle in WiT49 and 17.94 cells treated with DMSO, actD, BTZ, or combination actD + BTZ for 48 hours. (Student’s t-test for treated cells versus DMSO: * <0.05, **<0.01, ***0.001; G0,G1 Phase shown in green, and S Phase in blue). **(D)** Effect of 48-hour treatment WiT49 and 17.94 cells with DMSO, actD, BTZ, and combination actD + BTZ on levels of cell cycle and apoptosis markers by Western blot.

Our RPPA results (Fig. 1F) had shown a reduction in cell cycle markers in actD-treated WiT49, so we next examined how actD and BTZ affect cell cycle progression using flow cytometry (Figure 3C). Specifically, we treated WiT49 and 17.94 with actD and/or BTZ for 48 hours, and we measured the proportion of cells in each phase of the cell cycle using flow cytometry (Figure 3C). We found that actD increased the proportion of cells in G0/G1 and reduced the proportion in the S phase in both cell lines, though the difference was only statistically significant in WiT49. This is consistent with previous reports that low-dose actD treatment leads to G1 pause ^33,34^. In contrast, BTZ increased the proportion of cells in G2 and reduced the proportion in the S phase, consistent with previous reports that BTZ causes cells to accumulate in G2/M phase ^32^.

BTZ has previously been shown to trigger cell death, cell cycle arrest, and autophagy in cancer cell lines ^35–40^, so we next measured cell cycle and apoptosis markers using Western blots. In both cell lines, actD alone reduces cell cycle markers, including chromatin licensing and DNA replication factor 1 (CDT1), cyclin D2, cyclin E1, and cyclin A2 (Figure 3D) ^41–43^. These cell cycle regulators are synthesized and degraded in each turn of the cell cycle ^44,45^. These support our findings from CRISPR screen and protein arrays, which showed that actD induced a dependency on cell cycle genes and led to a fall in cell cycle markers (Figures 1E, 1F). In addition to impaired cell cycle progression, use of the proteasome inhibitor BTZ resulted in the accumulation of apoptotic markers in both cell lines (Figure 3D) ^46–49^. ActD alone induced a slight increase in cleaved caspase-7 and cleaved PARP in 17.94 cells, but both apoptosis markers were dramatically induced by BTZ in both cell lines. The combination of these effects supports the potential for using actD and BTZ to target anaplastic Wilms tumors.

### Combined treatment of actinomycin D and bortezomib in Wilms tumor xenografts suppresses growth *in vivo*

We next examined the effect of combining BTZ with actD against two anaplastic Wilms tumor lines *in vivo*: cell line-derived xenografts from 17.94 and the *TP53*-mutated patient-derived xenograft (PDX) line KT-53 ^50,51^. We implanted both lines into NSG mice and treated them with vehicle only, actD, BTZ, or both. We found that combination treatment significantly reduced tumor volume and conferred a significant survival advantage for mice bearing KT-53 xenografts compared to the other three arms (Figure 4A, 4B). Similarly, 17.94 xenografts treated with the combination of actD and BTZ were significantly smaller than those who received vehicle control or BTZ alone (Figure 4C, 4D). (Although it did not reach statistical significance, tumor volume in the combination treatment cohort was also slightly smaller than the actD-only arm.) In these treatments, BTZ was well tolerated and did not appear to add toxicity. There was no difference in body weight between mice receiving BTZ and vehicle (Supplementary Figure S6A, S6B). Similarly, the weights of mice treated with the combination of actD and BTZ was similar to the weights of mice treated with actD alone.

**Figure 4.**
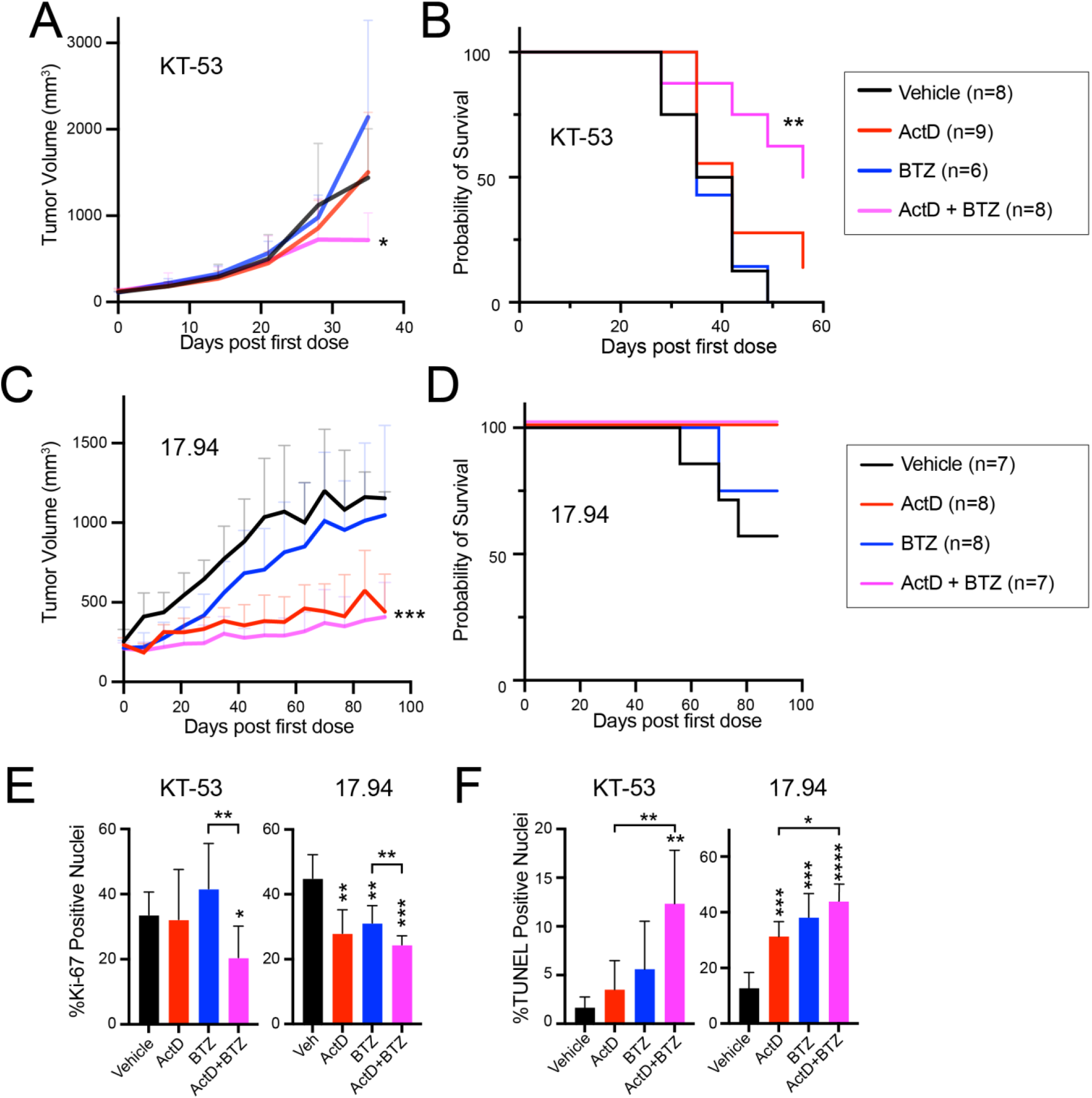
BTZ sensitizes subcutaneous anaplastic Wilms tumor xenografts to actD. **(A)** Subcutaneous tumor volumes of NSG mice bearing KT-53 xenografts treated with vehicle, actD, BTZ, and combination actD + BTZ (Student’s t-test of endpoint volumes of combination versus vehicle or actD only: * <0.05). **(B)** Kaplan-Meier survival curves for KT-53 tumor-bearing mice treated with vehicle, actD only, BTZ only, and combination (log-rank test versus vehicle ** <0.01). **(C)** Subcutaneous tumor volumes of NSG mice bearing 17.94 xenografts treated with vehicle, actD only, BTZ only, and combination (Student’s t-test of endpoint volumes of combination versus vehicle: ***<0.001). **(D)** Kaplan-Meier survival curves for 17.94 tumor-bearing mice treated with vehicle, actD only, BTZ only, and combination. **(E,F)** Quantification of Ki-67 positive nuclei **(E)** and TUNEL positive nuclei **(F)** in in KT-53 and 17.94 subcutaneous tumors treated with vehicle, actD only, BTZ only, or combination actD + BTZ (Student’s t-test p-value versus vehicle unless otherwise indicated: * <0.05, **<0.01, ***<0.001, ****<0.0001).

Based on the effects of actD and BTZ on cell cycle and apoptosis we had observed *in vitro*, we next measured cell cycle and apoptosis markers in these subcutaneous tumors. We used immunohistochemistry for Ki-67 as a marker of proliferation and terminal deoxynucleotidyl transferase dUTP nick-end labeling (TUNEL) assay as a marker of apoptosis. In both lines, tumors from combination-treated mice had a statistically significant reduction in proliferation compared to vehicle or BTZ alone (Figure 4E, Supplemental Figure S7A-S7D). Similarly, in both lines, combination treatment yielded significantly more apoptosis than vehicle and actD alone (Figure 4F, Supplemental Figure S7E, S7F). Taking these into consideration, we find that the compounding effects of adding BTZ to actD could be a powerful approach to targeting anaplastic Wilms tumor cells.

### Proteasome subunit expression levels correlate with outcome

Lastly, we examined expression of proteasome genes in publicly available RNA-seq data from 42 DAWT and 83 relapsed FHWT (rFHWT) samples generated by the NCI Therapeutically Applicable Research to Generate Effective Treatments (TARGET) project ^8^. Compared to rFHWT, the single most enriched KEGG gene set in DAWT was KEGG_PROTEASOME (Figure 5A, Supplementary Figure S8A, Supplementary Table S6). Similarly, two of the three most enriched Reactome gene sets were related to APC/C-mediated degradation of cell cycle proteins, suggesting that proteasomal degradation could promote proliferation in DAWT by degrading proteins in each cell cycle (Figure 5B, Supplementary Figure S8B, Supplementary Table S6).

**Figure 5.**
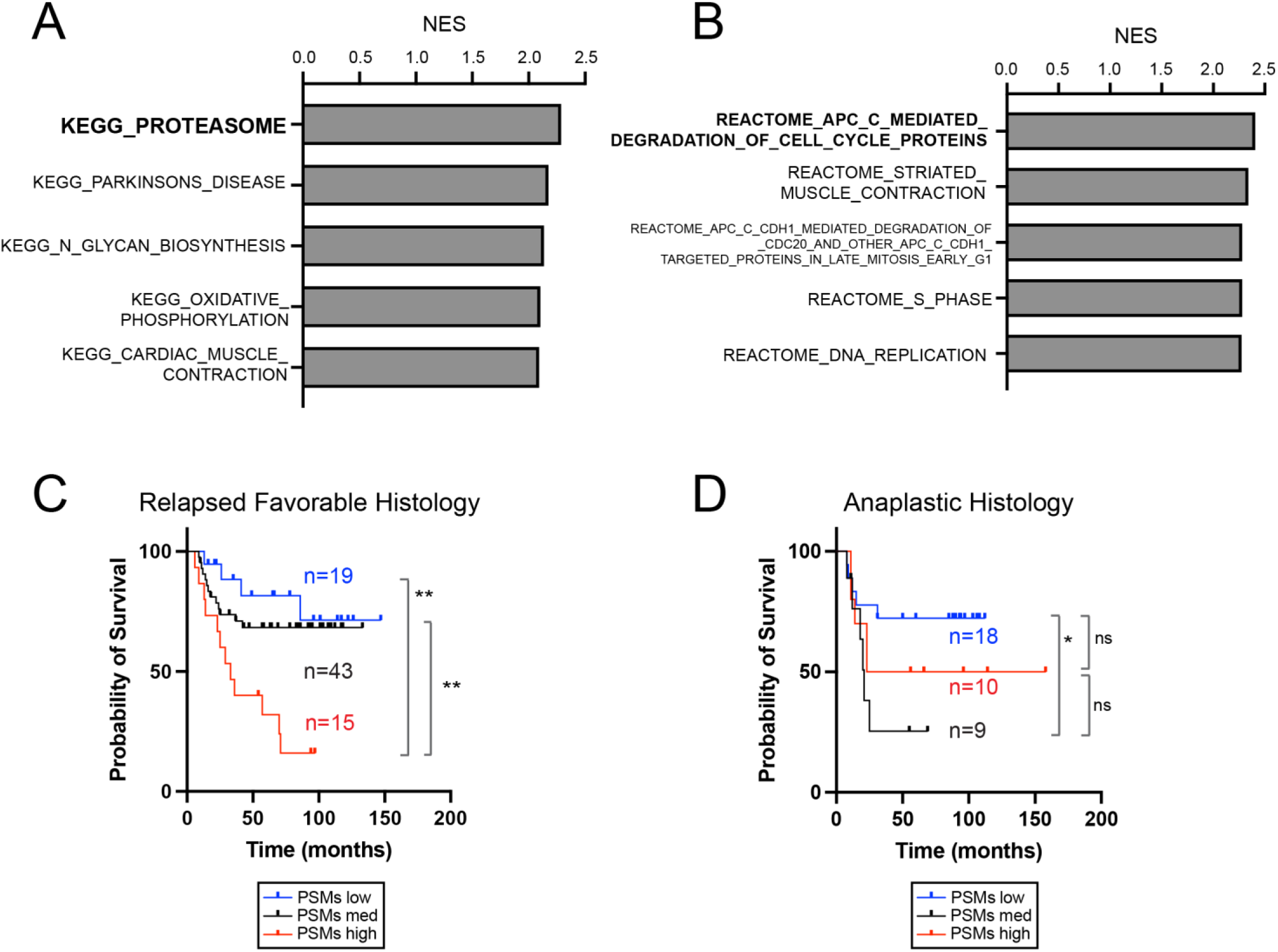
RNA expression of proteasome genes in TARGET anaplastic histology and relapsed favorable histology Wilms tumors. **(A)** Top KEGG pathways enriched in RNA-seq of DAWT versus rFHWT. **(B)** Top reactome pathways enriched in RNA-seq of DAWT versus rFHWT. **(C,D)** Overall survival of Wilms tumor patients stratified according to expression of proteasome enzymatic subunit genes PSMB1, PSMB2, PSMB5 among patients with relapsed favorable histology tumors **(C)** or anaplastic histology tumors **(D)**. (Log Rank test p-value: *<0.05, ** 0.01)

Next, we examined whether expression of proteasome subunits correlated with outcome in Wilms tumor. We stratified rFHWT and DAWT patients into three categories based on high, medium, or low expression of the enzymatic proteasome subunits *PSMB5* (β5), *PSMB6* (β1), and *PSMB7* (β2) based on expression z-scores for each gene. Tumors with at least one gene z-score ≥ +1 were categorized as “high”, while those with at least one gene z-score ≤ −1 were categorized as “low”. Samples with z-scores between −1 and +1 for all three genes were categorized as “medium”, while those with one gene z-score ≥ +1 and another gene z-score ≤ −1 were omitted.

Among rFHWT patients, proteasome-high patients fared far worse than proteasome-low or proteasome-medium patients (Figure 5C, Supplementary Figure S9A). However, this is a clinically heterogeneous population comprised of patients who would have received different treatments based on clinical stage, chromosomal loss-of-heterozygosity, and other factors. Thus, we next examined whether proteasome expression correlated with survival within each clinical stage. To increase statistical power, we combined the proteasome-low and proteasome-medium groups, which fared similarly to each other (Figure 5C). We then compared the survival of the proteasome-high group to the combined proteasome-low and proteasome-medium group within each stage (Supplementary Figures S9B-S9E). We found that high proteasome gene expression still correlated with significantly worse outcome in Stages II and IV (p=0.007 and 0.002, respectively). Although the correlation was not significant for the other groups, these analyses were limited by small sample sizes after subdividing by stage and proteasome expression. There was only one proteasome-high patient in the Stage I rFHWT cohort. In the Stage III rFHWT cohort, proteasome-high patients fared slightly worse but did not reach statistical significance (p=0.15).

Among DAWT patients, who do not usually receive actD-based therapy regimens, the relationship between proteasome expression and outcome was less evident; proteasome-medium patients fared worse than proteasome-low patients, but proteasome-high patients were not significantly different from proteasome-medium or -low ^8,52,53^ (Figure 5D). Together, these data suggest that proteasome subunit levels may underlie some of the clinical differences between DAWT and FHWT, and that high proteasome subunit levels correlate with poorer prognosis in rFHWT. Proteasome levels could be prognostic in some subgroups, and proteasome inhibition could benefit some of these patients.

## DISCUSSION

Developments in Wilms tumor research have illuminated the mutational, epigenetic, mRNA, and miRNA expression landscapes of these tumors, yet relevant targetable vulnerabilities remain elusive ^8,54–58^. Anaplastic Wilms tumors exhibit relative resistance to conventional chemotherapy, including actD, and little is known about how to overcome such resistance. Furthermore, despite its widespread use, little is known about how actD influences the translational landscape of cancer. Defining the mechanisms underlying these effects and their consequences could potentially uncover targetable vulnerabilities in relatively chemo-refractory anaplastic Wilms tumors, which could enhance outcomes while minimizing off-target toxicities in survivors. Through several orthogonal approaches, our work reveals that actD disrupts protein homeostasis in Wilms tumor and suggests proteasome inhibition as a potential targeted therapy for inducing actD sensitivity in anaplastic Wilms tumors.

Protein homeostasis involves a synchronized and responsive balance between protein synthesis and degradation, which are both affected by actD. Since protein synthesis is energy-intensive, particularly in rapidly proliferating cancers where proteins like cyclins are continuously synthesized and degraded, maintaining protein homeostasis is crucial to ensure optimal levels of amino acids and other nutrients for growth ^59–61^. Blocking protein synthesis with actD leads cells to upregulate Akt/mTORC1 signaling, which can increase the translation of ribosomal protein (RP) genes ^27,62–64^. Although we detected an increase in translational efficiency of RP genes, they did not accumulate by Western blot in our 72-hour treatments, possibly due to their instability without rRNA. Although the mTOR pathway is a potential therapeutic target in Wilms tumor and other cancers ^65–67^, the combination of actD and rapamycin did not consistently show synergy *in vitro* for either cell line, and rapamycin alone only exhibited a cytostatic effect. Our cells were in nutrient-rich media which may be compounding to the variables that dictate the effect of rapamycin ^66,68^, and our results do not rule out the possibility that other mTOR signaling inhibitors could still have potential.

The importance of protein homeostasis in Wilms tumor is also supported by other recent findings. Common Wilms tumor mutations interfere with protein homeostasis. For instance, recurrent mutations in *CTNNB1* and *MYCN* prolong their stability by interfering with their degradation by the proteasome ^8,69^. Other common mutations impair processing of microRNAs, which normally regulate translation of target transcripts ^54,57,58,70^. Recently, a small-molecule inhibitor of histone lysine demethylases KDM4A-C was found to act by reducing rRNA and RP transcripts in WiT49 cells, which led to a broad reduction in protein translation^71^. How actD or BTZ interacts with common Wilms tumor mutations or with KDM4A-C inhibition remains to be seen.

Based on our finding that proteasome subunits were preferentially translated after actD exposure, we found that proteasome inhibition with BTZ sensitizes cells to actD *in vitro* and *in vivo*. The proteasome is upregulated in response to proteotoxic stresses ^28,29,72^, and Akt/mTORC1 activates proteasome subunit expression via *Nrf1/NFE2L1* ^73–76^ to enhance the intracellular amino acid pool and the unfolded protein response. BTZ was the first proteasome inhibitor approved by the United States Food and Drug Administration ^77–79^, and it has been tested in pediatric cancer patients ^30,80,81^. However, other than a study showing a lack of cross-resistance between actD and BTZ in gliobastoma ^82^, no published studies have explored combinatorial treatment of actD and BTZ to our knowledge.

While proteasome inhibitors have been clinically successful in hematological cancers, single-agent proteasome inhibitor trials have not been as successful for solid tumors ^83,84^. It is thought that this could be due to insufficient pharmacokinetic distribution of BTZ. Strategies for enhancing the effect of BTZ in solid tumors include liposomal nanoformulations^85^ and combination with inhibiting other proteasomal components ^86^. Our data suggest that actD could be another way to enhance the effect of BTZ in solid tumors. While we did not observe increased toxicity with this approach in animals, it remains to be seen whether this combination would be tolerated in patients.

Compellingly, for our xenograft studies in both KT-53 and 17.94, systemic combination treatment reduced proliferative cells and increased apoptosis. While our *in vitro* experiments test sensitivity over a 72-hour period, *in vivo* experiments more closely model the once-weekly actD dosing used in patients^87^. Our study on KT-53 recapitulated a previously demonstrated insensitivity to actD ^51^, which we found could be overcome by BTZ. For 17.94, which was sensitive to nanomolar actD concentrations *in vitro*, actD alone was also effective *in vivo*, and adding BTZ produced a small additional effect. The different effects we observed in sensitivity to actD and BTZ suggest that other factors regulating protein homeostasis may also contribute to how cells respond to these drugs. Further studies involving more cell lines, other cancer types, or manipulation of individual factors that regulate protein homeostasis might help explain these differential responses. Together, our findings indicate that the combined use of actD and BTZ could be a promising strategy for combating therapy resistance in anaplastic Wilms tumors.

We demonstrated that actD impairs cell cycle progression *in vitro* and *in vivo*, which we attribute to dysregulated proteostasis ^15,88,89^. Normally, D-type cyclin levels accumulate in response to mitogenic signaling, promoting the transcription of S-phase genes such as E-type cyclins and chromatin licensing and DNA replication factor 1 (CDT1) ^44,90,91^. Consistent with our RPPA in WIT49, we observed in both cell lines that actD reduces Cyclin D2, Cyclin E1, and CDT1 levels, suggesting that actD prevents transition to S-phase, when cyclin A is produced. On the other hand, these proteins are normally targeted for proteasomal destruction by the SKP1-CUL1-F-boc protein (SCF) ubiquitin ligase ^92^, and indeed, we found that BTZ caused them to accumulate. Moreover, BTZ treatment induced apoptosis *in vitr*o and *in viv*o, consistent with studies on other cancer cell lines ^35–40^.

In the TARGET cohort ^8^, we found that higher levels of proteasome gene expression correlate with anaplastic histology and with poor outcome among some subgroups of relapsed favorable-histology Wilms tumors. Studies across other types of cancer show that the association of proteasome activity with prognosis is context-specific ^93^. Lower expression of proteasome subunit genes was associated with reduced survival in head and neck squamous cell carcinoma ^94^, while studies in breast cancer, glioma, and hepatocellular carcinoma found that higher expression of proteasome subunit genes is associated with worse survival ^95–97^. To our knowledge, this is the first study to correlate proteasome expression with worse outcome in Wilms tumor. Anaplastic Wilms tumors are strongly associated with *TP53* mutation ^7,8^, and *TP53* mutation is known to contribute to proteasome subunit overexpression ^98^. To our knowledge, this is the first study to explore the translational landscape of actD treatment in Wilms tumor. Through *in vitro* and *in vivo* models, we propose a model of protein homeostasis-dependent actD sensitivity (Supplementary Figure S10). Use of actD impairs ribosome biogenesis and cell cycle progression while upregulating proteasome activity. Higher proteasome capacity may help Wilms tumor cells escape front-line chemotherapeutics like actD, and this effect may be reversed with proteasome inhibition. Our complementary approaches converged on proteasome activity as a potential mechanism for chemoresistance. Repurposing widely used drugs like BTZ could allow for an accelerated path to clinical translation. More broadly, this strategy could improve treatment not only for Wilms tumor, but also for other tumors where actD is used, such as rhabdomyosarcoma and Ewing sarcoma.

## Limitations of this study

Our conclusions are limited by the fact that our *in vitro* studies focused on two anaplastic Wilms tumor cell lines, and our *in vivo* studies focused on two anaplastic Wilms tumor PDX lines. We did not compare them to favorable histology Wilms tumor cell lines or xenografts, and we cannot know how our observations might extend to non-anaplastic Wilms tumors. Similarly, our analysis of human Wilms tumor RNA-seq was based on published TARGET data, which only includes rFHWT and DAWT. As such, we do not know whether expression of proteasome components would predict outcome in a population of FHWT at diagnosis. Similarly, as these are retrospective data, the predictive power of these expression patterns needs to be validated in an independent cohort. Lastly, just as *in vitro* findings may vary from *in vivo* results, pre-clinical models cannot replace clinical trials. To our knowledge, actD and BTZ have not been given in combination, and the toxicities of such a regimen are unknown.

## ACKNOWLEDGMENTS

This work was supported by funds from the Pablove Foundation (to P.D.B.T.); Alex’s Lemonade Stand Foundation (to K.S.C.); Cancer Prevention and Research Institute of Texas (RP180805 to L.X., RR180071 to K.S.C.); US Department of Defense (KC220019 to K.S.C.); and National Cancer Institute (K08CA207849 to K.S.C., R21CA259771 and P30CA142543 to L.X., and Cancer Center Support Grant P30CA142543). This research used computational resources provided by the BioHPC supercomputing facility in the Lyda Hill Department of Bioinformatics at UT Southwestern, which is supported by Cancer Prevention and Research Institute of Texas (RP150596). The Functional Proteomics Reverse Phase Protein Array Core was supported in part by The University of Texas MD Anderson Cancer Center, P30CA016672 and R50CA221675. The authors also wish to thank Dr. Philipp Scherer of UT Southwestern Medical Center for use of equipment, Lindsay Mendyka, Lane Beeman, and Chelcea Morris for their technical assistance.

## STAR METHODS

### Experimental Model and Study Details

#### Tissue culture

The anaplastic, *TP53*-mutated stage IV Wilms tumor lung metastasis cell line WiT49 (RRID: CVCL_0583) from a female patient was maintained in Dulbecco’s Modified Eagle Medium (DMEM) with 1000 mg/L glucose, 4 mM L-glutamine, 1 mM pyruvate (Gibco 11995065) supplemented with antibiotic-antimycotic (Gibco 15240062) and fetal bovine serum (Sigma F2442) to 10% final concentration in 37°C at 5% CO_2_. A second anaplastic, *TP53*-mutated Wilms tumor cell line 17.94 (Ximbio 153333, RRID:CVCL_D704) originally collected from a female patient was grown in DMEM supplemented with antibiotic-antimycotic and heat inactivated fetal bovine serum (Gibco 16140063) to 20% final concentration in 37°C at 5% CO_2_. Cell line identity is annually verified by short tandem repeat (STR) genotyping (last verified November 2024). They are also tested biannually for mycoplasma contamination (Latest negative screen November 2024, Lonza LT07-318).

#### Xenograft tumor models

All experiments involving animals were reviewed, approved, and monitored for compliance by the UT Southwestern Institutional Animal Care and Use Committee.

To generate xenografts, log growth-phase 17.94 cells or freshly-thawed cryopreserved *TP53* mutated anaplastic Wilms tumor PDX line from a female patient KT-53^51^ were injected subcutaneously into the flanks of NOD *scid* gamma (NSG) mice (8-12 weeks at transplantation) in 1 part DPBS and 1 part Matrigel (Corning 354234) at 4 million 17.94 per 100 µl or 1.6 million KT-53 cells per 100 µl per mouse. The mice were then randomized into four anticipated treatment arms, with each sex represented to within n±2 of each other in each treatment group. Tumor growth was measured once a week with calipers in two dimensions. Tumor volume was calculated by dividing the product of the length and the square of the width by 2. Upon reaching the thresholds of 150 mm^3^ for 17.94 and 100 mm^3^ for KT-53, mice were began one of four treatments: actD only, BTZ only, both actD + BTZ and vehicle. Mice in the actD only and actD+BTZ arms were dosed with actD (in 2.5% DMSO, 40% PEG300, 5% Tween-80, 52.5% saline) once a week intraperitoneally (0.2 mg/kg for 17.94, 0.15 mg/kg for KT-53). Mice in the BTZ only or actD+BTZ arms were dosed with BTZ (in 0.5% DMSO, 30% PEG300, 69.5% distilled deionized water) once a week intravenously (0.2mg/kg). Mice in the vehicle arm received both solvents once a week. Treatment continued until tumor volumes reached 1,500 mm^3^ or at maximum 8 weeks for KT-53 and 12 weeks for 17.94 post first dose—conditions at which all but actD+BTZ, or DMSO and BTZ respectively are either already terminated or at least 1,000 mm^3^. At the end of therapy, tumors were harvested for histological processing. Statistical analyses for tumor volume were performed by two-tailed Student’s T-test for each pairwise comparison between the four treatment arms. Kaplan-Meier survival curves were compared between each group by log-rank test.

### Method Details

#### Ribosomal RNA quantification

To determine the effect of actD on rRNA expression, WiT49 or 17.94 cells were seeded at ∼50% confluency in 3 different plates. The following day, cells were treated with either vehicle (DMSO) or 2nM actD for 6 hours or 24 hours. At experiment endpoint, total RNA was extracted using the miRNeasy kit with DNase I digestion (Qiagen 217004 and 79254), and cDNA synthesis was performed with iScript™ Reverse Transcription Supermix (Bio-Rad 1708841). Quantitative PCR (qPCR) was performed in four technical replicates per condition using iTaq™ Universal SYBR® Green Supermix (Bio-Rad 1725125) with primers listed: *18s* rRNA (GTAACCCGTTGAACCCCATT, CCATCCAATCGGTAGTAGCG); *45s* pre-rRNA (ACCCACCCTCGGTGAGA, CAAGGCACGCCTCTCAGAT); *GAPDH* (CGGAGTCAACGGATTTGGT, ACCAGAGTTAAAAGCAGCCC). Relative expression was calculated by the 2^-ΔΔCt^ method by normalizing each rRNA species to *GAPDH*. Significance was determined by unpaired two-tailed Student’s T-test versus vehicle control. A second biological replicate was performed as above to confirm the results.

#### Protein synthesis quantification

To determine the effect of actD on protein synthesis using Click-iT® AHA (L-azidohomoalanine, Invitrogen C10102), WiT49 or 17.94 cells were first seeded at 30% confluency in 15 cm plates and treated the following day with 2nM actD or DMSO in complete media. Following 48 hours of drug exposure, the media was replaced with methionine-free complete media [DMEM without methionine (Gibco 21013024), with 10% dialyzed FBS (Gibco A3382001), 1x antibiotic-antimycotic (Gibco 15240062), 0.2 mM L-Cystine (Thermo Scientific J61651.09), 4mM L-Glutamine (Gibco 25030149), 1mM Sodium Pyruvate (Gibco 11360070)] containing DMSO or 2nM actD for 30 minutes to wash out any residual methionine. After 30 minutes in methionine-free media, the media was supplemented with Click-iT® AHA to a final concentration of 50µM, and the cells were returned to the incubator for 4 hours. Whole protein lysates were extracted from these cells with RIPA buffer (Sigma R0278), supplemented with protease and phosphatase inhibitors (Invitrogen A32961), sonicated, and quantified by BCA protein assay (Thermo Scientific 23227). For AHA-detection, we used the Click-&-Go® Click Chemistry Reaction Buffer Kit (Vector Laboratories Inc. CCT-1001) with AZDye 680 Alkyne (Vector Laboratories Inc. CCT-1001) according to manufacturer recommendations. For each condition, 20µg of total protein lysates were reacted with the alkyne reagent at a final concentration of 20µM. Following denaturing SDS-PAGE, the gel was imaged (LI-COR Biotech) to detect proteins with AZDye 680 conjugated AHA. Results were performed with biological replicates to confirm reproducibility.

Additionally, we detected nascent protein synthesis using O-propargyl-puromycin (OPP) incorporation. WiT49 cells were seeded at ∼30% confluency in cell culture chamber slides. The following day, cells were treated with either vehicle (DMSO) or 2nM actD for 48 hours or 72 hours. At the endpoint, protein synthesis was measured using a fluorescence-based O-propargyl-puromycin (OPP) incorporation kit (Invitrogen C10457) according to manufacturer protocols. Nuclei were counterstained with DAPI (4’,6-diamidino-2-phenylindole). Each condition was photographed at 5 non-overlapping regions at 20x magnification on a BZ-X810 Fluorescence microscope (Keyence Corporation) and quantified using Fiji ^99^. Necrotic regions were disregarded. Relative fluorescence units were calculated from the blue (DAPI) and red (OPP-Alexa Fluor 488) channels. The number of nuclei was counted from the DAPI channel using the Watershed and Analyze Particles functions in Fiji. Raw protein synthesis was calculated as the average red fluorescence intensity in pixels with any fluorescence using the Area and Integrated Density functions in Fiji. For each image, raw protein synthesis was then normalized to the number of nuclei in that field of view. Each field of view was treated as a technical replicate, and statistical analysis was performed by unpaired two-tailed Student’s t-test against untreated cells.

#### Reverse phase protein array

WiT49 cells at 20% density were treated with 1 nM actD or DMSO as vehicle control for 72 hours in duplicate. 1×10^6^ cells were collected, washed, and frozen in liquid nitrogen, and sent to the MD Anderson Functional Proteomic Reverse Phase Protein Array Core ^100–102^. Normalized, log2-transformed, median-centered values from “validated” antibodies were used for analysis ^103^. The log2-fold-change for each antibody was calculated as the difference between the means of actD-treated and DMSO-treated samples, and the standard deviation for each antibody was calculated as the standard deviation of all four values.

#### Ribosome profiling

WiT49 cells were incubated in complete growth medium with 2nM actD or DMSO for 6 or 72 hours, in three technical replicate plates per condition. For 2-week long treatments, WiT49 were maintained and passaged in two 7-day cycles of drug/vehicle-supplemented media for 3 days, followed by drug-free complete growth media for 4 days, three technical replicate plates per condition.

1.5 × 10^7^ cells were used for each replicate for ribosome sequencing. Ribosome footprints were isolated based on previous publications ^104,105^, and each RNA sample was spiked with 24 fmol of the synthetic 28-nt RNA oligo 5’-AUGUAACACGGAGUCGACCCGCAACGCGA-3’ ^106^. rRNA depletion was performed using the Low Input RiboMinus™ Eukaryote System v2 (Thermo Fisher A15027), and sequencing libraries were generated using the NEBNext® Small RNA Library Prep Set for Illumina® (NEB E7330). Libraries were pooled and sequenced using NextSeq 500 High Output, single-end reads, 75 cycles. In parallel, RNA was extracted for total RNA sequencing using the Direct-zol RNA Miniprep Kit (ZymoResearch R2050) or the miRneasy Mini Kit (Qiagen 217004) according to manufacturer protocols, followed by rRNA depletion as above and DNase-treatment. Sequencing libraries were prepared with the TruSeq Stranded Total RNA-seq Sample Prep Kit from Illumina and sequenced paired-end for 6-hour and 72-hr, and single-end for 14-day time points.

Trimmed reads from ribosome profiling and RNA sequencing were aligned to the hg38 reference genome using HISAT2 ^107^. For ribosome profiling, reads longer than 30 nucleotides were filtered out, and remaining reads were assembled to GENCODE v26 transcript annotations using StringTie ^108^. Ribosome profiling quantifications were normalized to mapped spike-in reads, and RNA-seq reads were normalized to million mapped reads. Translational efficiency (TE) for each gene was calculated by dividing the number of normalized ribosome footprint reads by the number of normalized RNA sequencing reads. Differential TE was calculated using t-tests on log2-transformed TE of three actD-treated technical replicates versus three DMSO-treated technical replicates. Geneset enrichment analysis (GSEA) was performed using the fgsea package (v1.26.0) on log2-fold change of TE using gene sets from MSigDB v7.1 ^17,109–111^.

#### CRISPR screen

The CRISPR knock-out screen was performed on WiT49 cells using the Human Brunello CRISPR knockout pooled library, a gift from David Root and John Doench (Addgene 73178). The plasmid library was propagated, verified for maintenance of representation, and transfected as recommended ^112^. The viral supernatant was transduced into 1.0 × 10^8^ WiT49 cells, at multiplicity of infection of 0.3. Forty-eight hours after transduction, the cells were selected with 0.5 µg/ml puromycin for 7 days. After selection, 3 × 10^7^ transduced cells were collected as transduction reference, and the rest were split into four dishes, two per treatment condition. These remaining cells were expanded and treated with either 2nM actD or DMSO for 4 days. The media was then replaced in both conditions for a washout of 3 days without actD or DMSO. Following this, another 7-day cycle (4 days actD or DMSO treatment followed by 3 days washout) was carried out. Finally, genomic DNA was extracted using DNeasy Blood & Tissue Kit (Qiagen 69506), and a sequencing library was prepared from genomically-integrated single guide RNA (sgRNA) sequences by PCR using barcoding and staggered primer pairs as in Supplementary Table S7 ^112^. These were gel-size selected and sequenced at ∼2.4 million single-end reads each, 100-bp length.

Computational analysis of CRISPR screens was performed using the MAGeCKFlute pipeline following published guidelines ^113^. In brief, we mapped reads using the command ‘mageck count’, and we tested sgRNA knockout efficiency using the command ‘mageck mle’. Downstream analysis was performed using the FluteRRA function in the MAGeCKFlute R package, with the ‘mageck mle’ output file as the input. To perform KEGG pathway enrichment analysis, the ‘mageck pathway’ command was used with the KEGG gene set from the Molecular Signatures Database (v7.1) ^109^. ^113,114^

#### Western blot

For the actD timecourse, WiT49 or 17.94 was seeded at ∼30% confluency overnight. The following day, without replacing the media, actD (or an equivalent volume of DMSO) was spiked into the media to a final concentration of 2 nM, for 72-hour treatment. The next day, another plate of WiT49 or 17.94 began 48-hour treatment with 2nM actD. We repeated this the following day for the 24-hour treatment and 18-hour treatments. Upon completion of the allotted incubation times, we collected cells for protein lysate extraction. For actD and BTZ *in vitro* drug treatments, we seeded WiT49 or 17.94 at 20-30% confluency overnight. We then treated cells with either 2nM actD alone, 8nM BTZ alone, 2nM actD + 8nM BTZ, or DMSO at equivalent volume without changing media. After 48 hours of drug exposure, cells were collected for protein lysate extraction.

Protein lysates were extracted from flash-frozen pellets with RIPA buffer (Sigma R0278), supplemented with protease and phosphatase inhibitors (Invitrogen A32961), sonicated, and quantified by BCA protein assay (Thermo Scientific 23227). Following denaturing SDS-PAGE and transfer to PVDF or nitrocellulose membranes, blots were blocked with 5% bovine serum albumin in tris-buffered saline with Tween-20. Primary antibodies used are as follows and were diluted to 1:3,000 unless otherwise stated: Tubulin (Cell Signaling Technology 3873, RRID:AB_1904178), GAPDH (Cell Signaling Technology 97166, RRID:AB_2756824), Caspase-7 (Cell Signaling Technology 12827, RRID:AB_2687912), cleaved Caspase-7 (Cell Signaling Technology 8438, RRID:AB_11178377; 1:1,000), Cyclin D2 (Cell Signaling Technology 3741; RRID:AB_2070685), CDT1 (Cell Signaling Technology 8064, RRID:AB_10896851), Cyclin E1 Cell Signaling Technology 20808, RRID:AB_2783554), Cyclin A2 (Cell Signaling Technology 91500RRID:AB_3096041), PARP (Cell Signaling Technology 9532, RRID:AB_659884), cleaved PARP (Cell Signaling Technology 5625, RRID:AB_10699459; 1:1,000), POMP (Cell Signaling Technology 15141, RRID:AB_2798726), PSMA6 (Cell Signaling Technology 2459, RRID:AB_2268879), PSMB1 (Invitrogen PA5-49648, RRID: AB_2635102; 1:5,000), PSMB2 (Proteintech 15154-1-AP, RRID:AB_2300322; 1:5,000), PSMB5 (Cell Signaling Technology 12919, RRID:AB_2798061; 1:5,000), PSMD1 (Sigma SAB2104781, RRID:AB_10668741), RPL5 (Cell Signaling Techonology 51345, RRID:AB_2799391), RPL7 (abcam ab72550, RRID:AB_1270391), RPL11 (Cell Signaling Technology 18163, RRID:AB_2798794), RPL26 (Cell Signaling Technology 2065, RRID:AB_2146242), Total AKT (Cell Signaling Technology 9272, RRID:AB_329827), Phospho-AKT (Ser473) (Cell Signaling Technology 4060, RRID:AB_2315049; 1:1,000), Total 4E-BP1 (Cell Signaling Technology 9452, RRID:AB_331692), Phospho-4E-BP1 (Thr37/46) (Cell Signaling Technology 2855, RRID:AB_560835; 1:1,000), Total P70-S6K1 (Cell Signaling Technology 34475, RRID:AB_2943679), and Phospho-P70-S6K1 (Thr389) (Invitrogen 710095, AB_2532559; 1:1,000). HRP-conjugated secondary antibodies used are Anti-Mouse IgG (Cell Signaling Technology 7076, RRID:AB_330924; 1:10,000) or Anti-Rabbit IgG (Cell Signaling Technology 7074, RRID:AB_2099233; 1:5,000). Each western blot run was performed at minimum with a second biological replicate, and band intensities were quantified using Fiji ^115^.

#### In vitro drug inhibition

To quantify drug inhibitory activity, 1,000 cells per well were seeded in black-walled 96-well plates at 100 µl/well. The following day, serial dilutions of actD (Sigma A1410), BTZ (Sigma 5043140001), or Rapamycin (Selleckchem S1039) were added in 3 technical replicate wells per dose. Vehicle control was also included to yield equivalent final concentrations of DMSO (<0.1%) in all wells. For synergy determination experiments, each drug of interest was serially diluted alone and then mixed into a range of combinations. Each dose was then added onto the cells in triplicate. After 72 hours, cell density was assayed by adding 20 µL 0.15 mg/ml resazurin in phosphate-buffered saline in each well, including cell-free wells as background controls and drug-free wells as normalization controls. After 1-4 hours at 37°C, fluorescence was measured using a microplate reader with excitation set at 550 nm and emission at 590 nm. Combination treatment effects were determined using the Loewe synergy model dose-response calculation and 95% confidence interval determination through Synergyfinder+ ^116^. Each experimental condition with three technical replicates was repeated at least twice to confirm reproducibility of outcomes.

#### Proteasome activity assay

To measure proteasome enzymatic activity, we seeded 1,000 cells per well concurrently in duplicate 96-well plates: one black-walled (for quantifying cell density) and one white-walled (for quantifying proteasome activity). The following day, cells were treated with each of the following conditions in quadruplicate with identical conditions in both plates: 2nM actD, 2 nM actD + 1 mM MG-132 (Selleckchem S2619), 8 nM BTZ, 8 nM + 1 mM MG-132, DMSO, or DMSO + 1 mM MG-132. Each condition was performed with or without MG-132 to correct for the level of non-proteasome chymotrypsin-like activity. Wells without cells were used as background controls. After 48 hours, the black-walled plate was used to determine cell viability using resazurin as described above, and the white-walled plate was used to quantify proteasome enzymatic activity. We used Proteasome-Glo™ Chymotrypsin-like Assay (Promega G8621) and calculated the proteasome enzymatic activity in accordance with manufacturer recommendations. Briefly, after subtracting background controls, we normalized each condition to viability based on the corresponding wells in the black-walled plate to account for differences in cell density. We then subtracted the normalized luminescence of each condition with MG-132 from the corresponding wells without MG-132 to yield the viability-normalized luminescence attributable to proteasome activity. Statistical significance was calculated between treatment conditions using unpaired, two-tailed Student’s T-test. The assay repeated at least twice for both cell lines to confirm reproducibility of outcomes.

#### Cell Cycle Phase Determination

To characterize the cell cycle phase distribution of WiT49 or 17.94 following drug treatment, we used flow cytometry with propidium iodide (Invitrogen F10797) with EdU (Invitrogen C10636) according to manufacturer recommendations. In detail, WiT49 or 17.94 cells were seeded at 30% confluency in 6-well plates. To perform cell cycle synchronization, the media in all but one (asynchronous control) well was replaced with DMEM (Gibco 11995065) with antibiotic-antimycotic (Gibco 15240062) and fetal bovine serum (Sigma F2442) at a reduced concentration of 0.2%. After a 24-hour incubation at 37°C at 5% CO_2_, the media was replaced with complete media with the full concentration of serum (10% for WiT49 and 20% for 17.94), with either 2nM actD, 8nM BTZ, 2nM actD + 8nM BTZ, or DMSO. After 48 hours of drug exposure, EdU (Invitrogen C10636) was added to each well, to a final concentration of 10µM, for 2 hours. The cells were gently dissociated, washed, fixed, and permeabilized in preparation for flow cytometry analysis according to the manufacturer protocols. The stained cells were resuspended in propidium iodide staining solution (Invitrogen F10797) for at least 15 minutes at room temperature and were analyzed by flow cytometry BD FACSCAnto (BD Life Sciences). The results were analyzed using FlowJo (BD Life Sciences). Populations in each cell cycle phase were obtained in triplicate runs and compared by Student’s T-test against the cell cycle phase proportions of the DMSO condition.

#### IHC & TUNEL

Two representative samples from each treatment arm were formalin-fixed, paraffin-embedded, and sectioned at 4 µm thickness. These were used for immunohistochemistry (IHC) and TUNEL assay.

For IHC, antibodies used were Ki-67 (Abcam ab15580, RRID:AB_443209, 1:1,000 for primary) with anti-Rabbit HRP (Vector Laboratories Inc. MP-7451, RRID:AB_2631198 for secondary). Each slide was photographed in at least four distinct 40x magnified fields of view (Keyence Corporation). Numbers of total nuclei and DAB-positive nuclei in each field were quantified using Fiji ^115^. Non-tumor regions were omitted from quantification. Statistical analysis was performed by unpaired two-tailed Student’s t-test between each pair of treatment arms.

To measure apoptosis in tumor sections, we used Click-iT™ Plus TUNEL Assay kit (Invitrogen C10617) according to the manufacturer recommendations and counterstained nuclei with DAPI. Each tissue slide was photographed in at least four distinct 40x magnified fields of view (Keyence Corporation). Numbers of total nuclei and TUNEL-positive nuclei and quantified using Fiji ^115^. Non-tumor regions were omitted from quantification. Statistical analysis was performed by unpaired two-tailed Student’s t-test between each pair of treatment arms.

#### TARGET database reanalysis

Tables of Wilms tumor RNA-seq counts and clinical annotation were downloaded from the NCI TARGET^8^ website on May 21, 2019. Protein-coding genes were identified based on ENSEMBL v86 annotations. Differential expression analysis for DAWT versus FHWT was performed with DESeq2 (v1.40.2), followed by gene set enrichment analysis using fgsea (v1.26.0) based on MSigDB v7.1 gene sets ^17,111,117^.

We compared outcomes for these patients according to expression of proteasome enzymatic subunits in cBioportal ^52,53^. Specifically, we obtained RNA-seq RPKM z-scores for *PSMB5*, *PSMB6*, and *PSMB7,* compared to tumors that are diploid for that gene. For patients with multiple samples, the primary tumor sample was used. Log-rank tests were used to compare survival curves.

#### Data availability

Ribosome profiling and accompanying RNA-seq data from WiT49 are deposited in the NCBI GEO database under accession GSE270330.

### Statistical Analysis

Statistical analyses, standard error, or 95% confidence intervals for the above were performed with the aid of GraphPad Prism software unless otherwise indicated, and details of each test used are referenced in figure legends. All data with replicates are presented as mean and error bars shown are for standard error of the mean. Values of p < 0.05 are considered statistically significant.

## SUPPLEMENTARY FIGURES

**Supplementary Figure S1.**
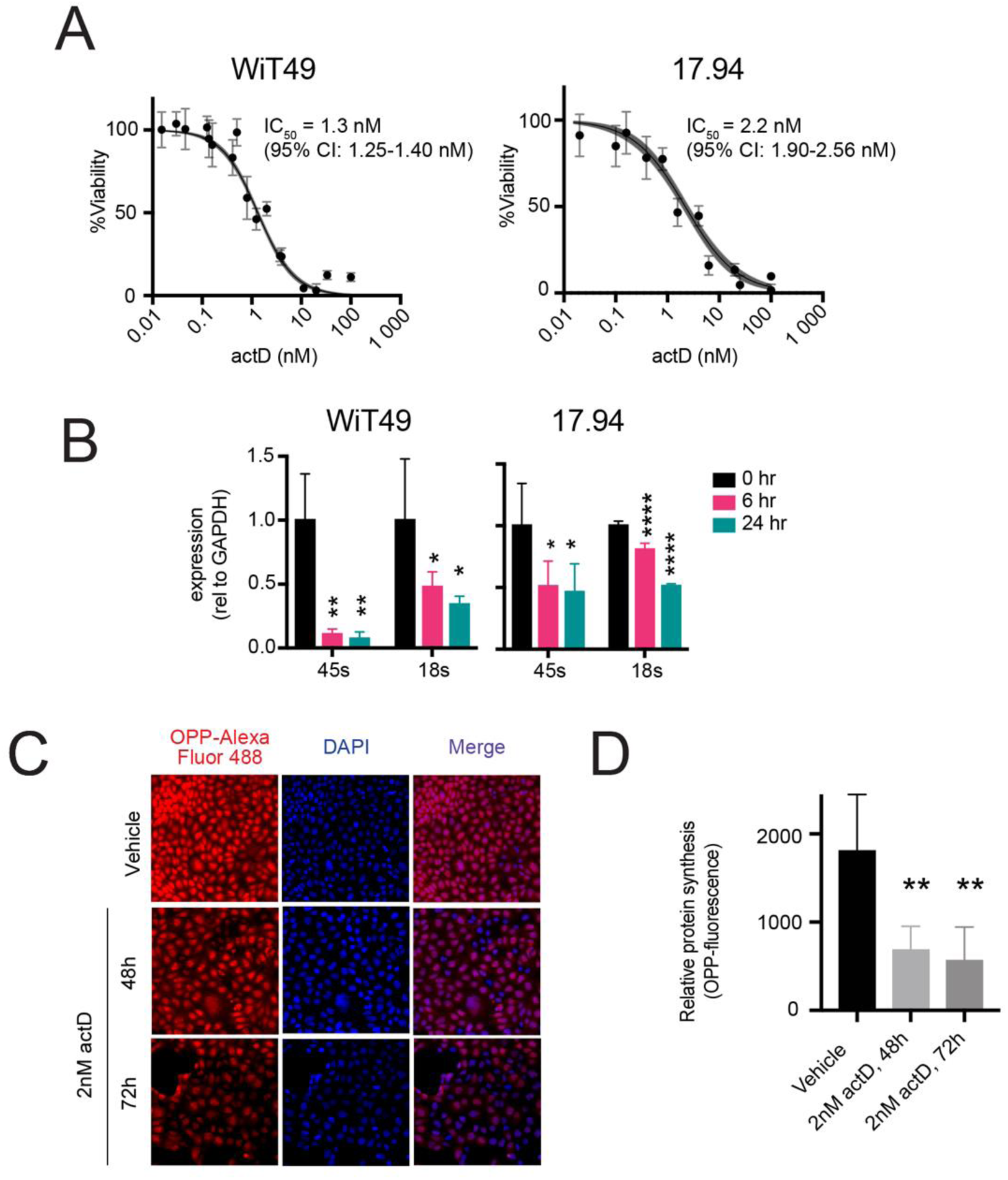
ActD alters global translation in WiT49 anaplastic Wilms tumor cells. **(A)** ActD kill curves for anaplastic Wilms Tumor cell lines WiT49 and 17.94 with IC50 values indicated (four-point regression line with shaded region representing 95% confidence interval). **(B)** Levels of 45s pre-rRNA and 18s mature rRNA in anaplastic Wilms tumor cell lines WiT49 and 17.94, quantified by qPCR following actD treatment for 6 and 24 hours. **(C,D)** Micrographs **(C)** and quantification **(D)** of protein synthesis assay. Red fluorescence measures O-propargyl-puromycin (OPP) incorporation relative to total area of WiT49 cells treated with 48 and 72 hours of 2nM actD or vehicle DMSO. (Student’s t-test vs. vehicle, ** p<0.01.)

**Supplementary Figure S2.**
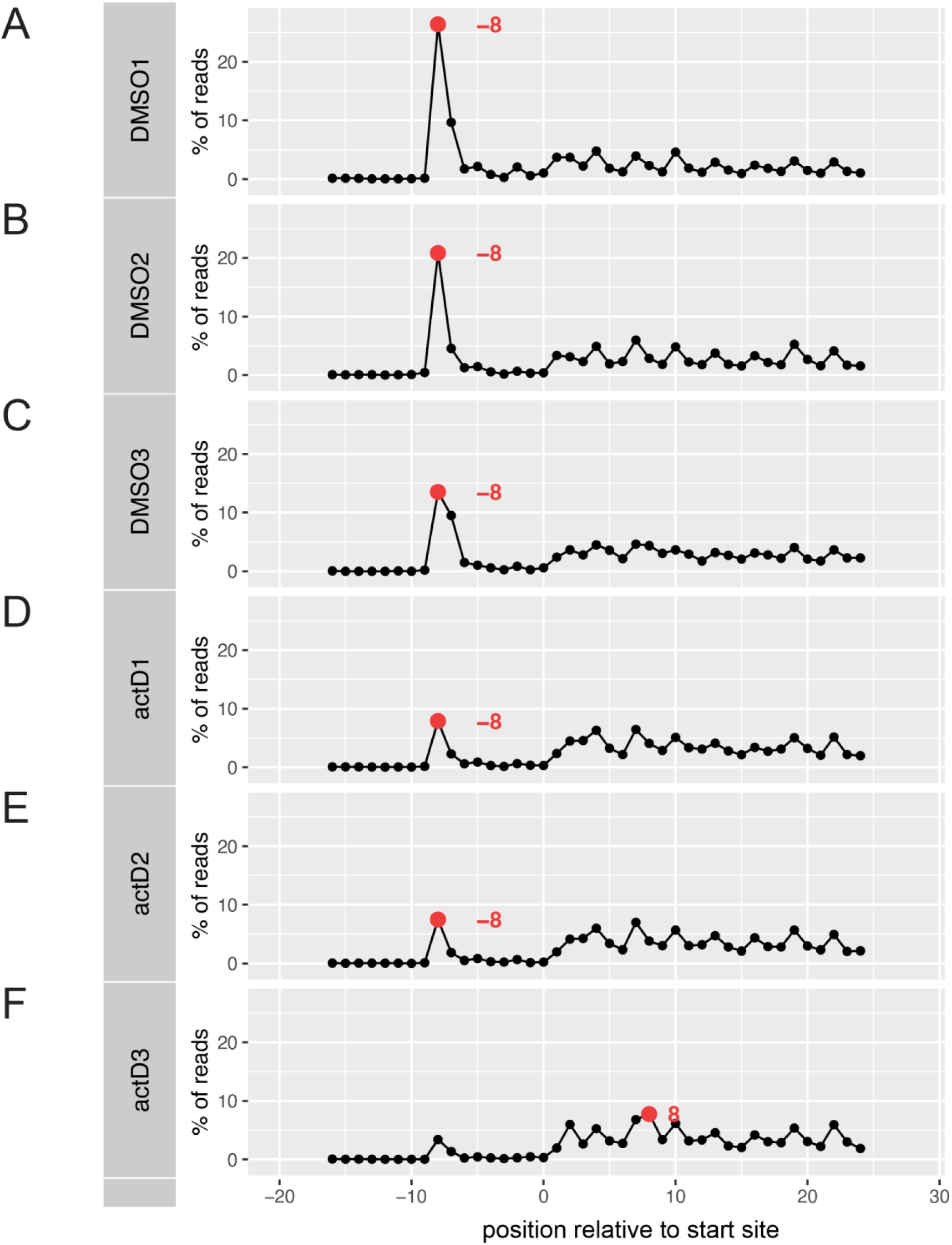
Ribosome footprints are depleted at translation initiation sites in actD-treated WiT49. **(A-F)** Proportion of left edge of ribosome footprints (y-axis) relative to translation initiation site (0 on x-axis), on the top 30% most abundant transcripts, in samples treated with DMSO **(A-C)** or actD **(D-F)** at 72 hours. Peak position highlighted in red.

**Supplementary Figure S3.**
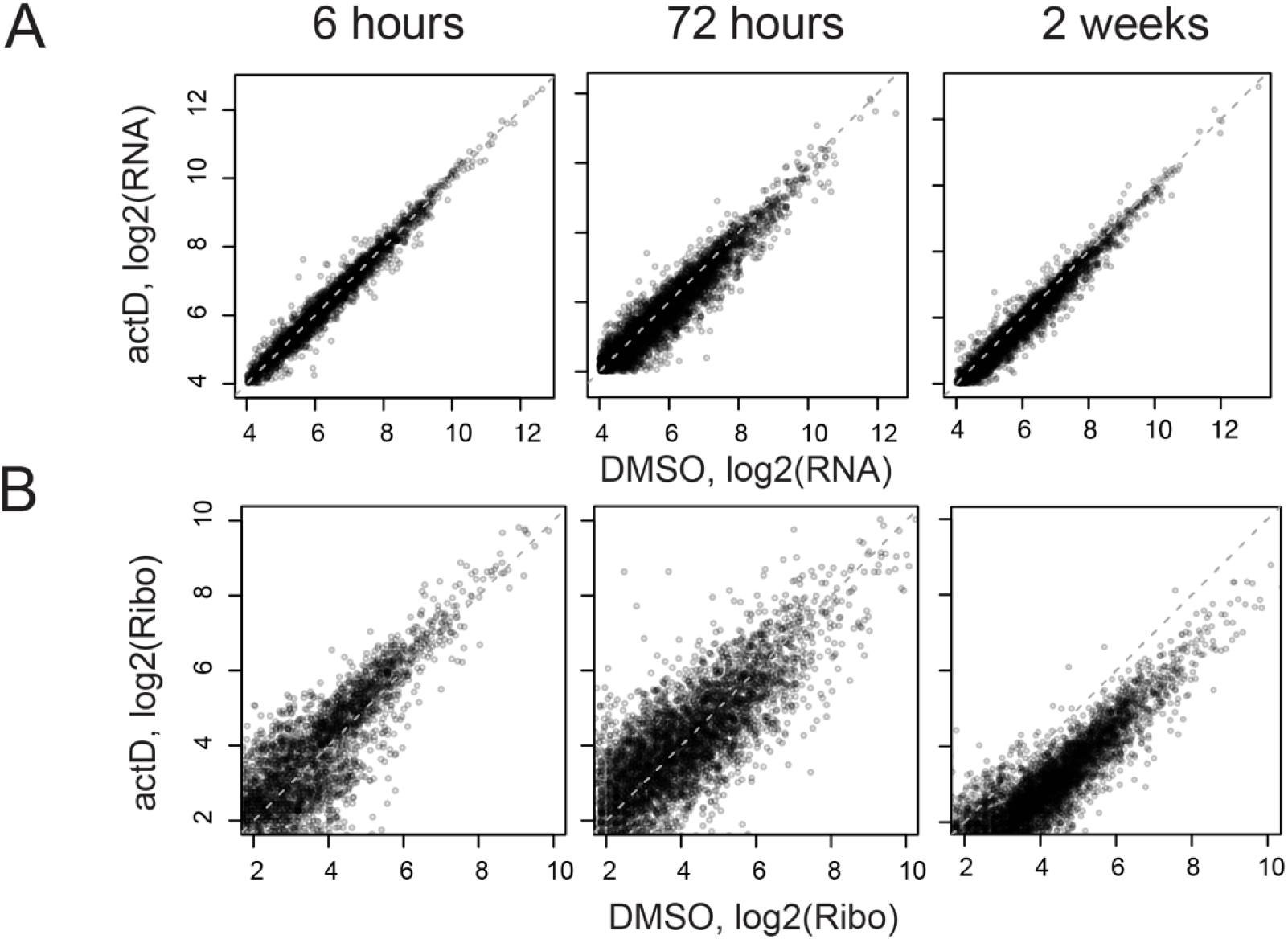
Ribosome profiling metrics in actD-treated WiT49. **(A,B)** Log2-transformed counts aligning to each gene, by RNA-seq **(A)** and translation efficiency by ribosome profiling **(B)** after 6-hour, 72-hour, and 2-week treatments of WiT49 with actD (y-axes) versus DMSO (x-axes).

**Supplementary Figure S4.**
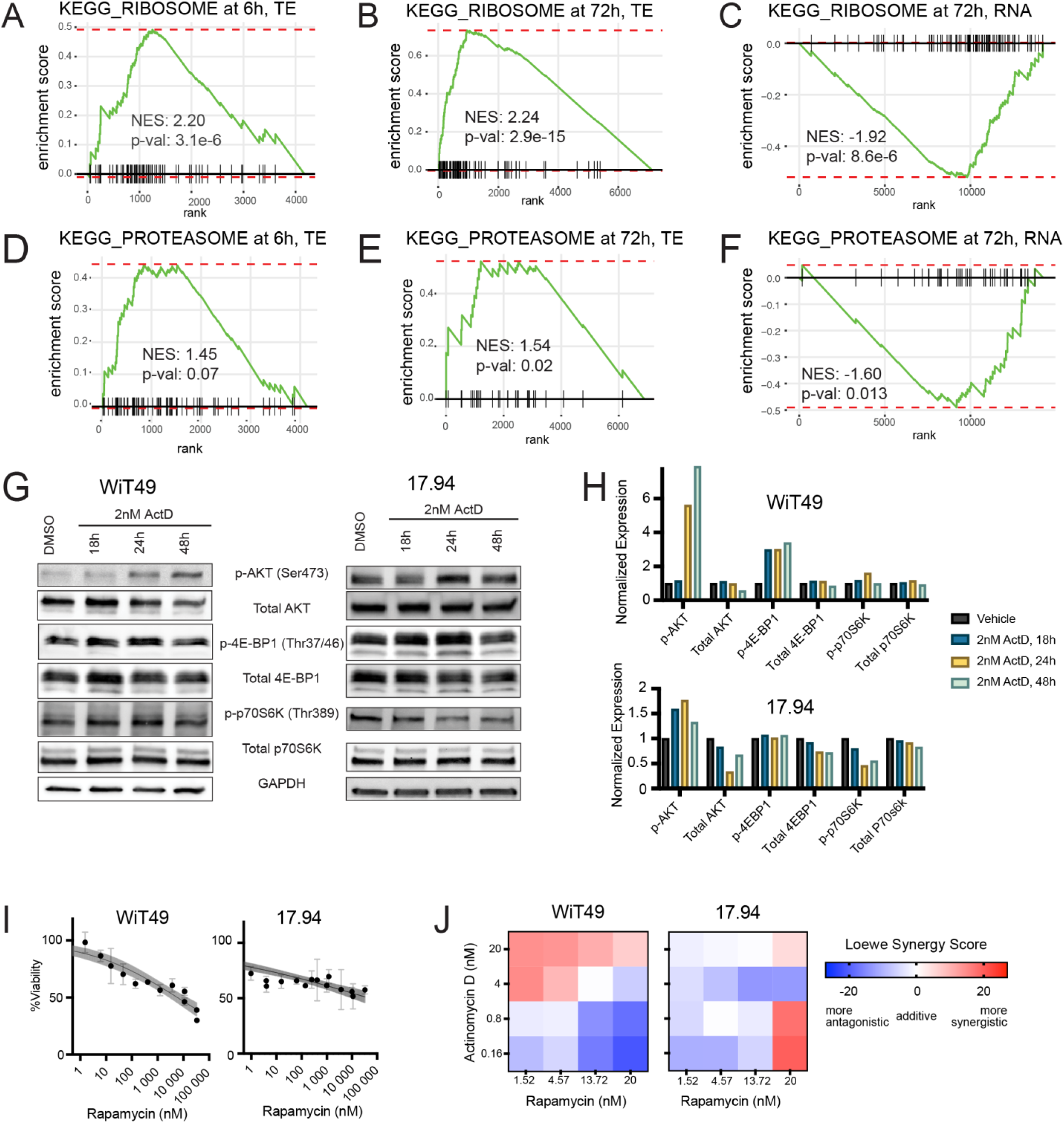
ActD induces translation of ribosome and proteasome subunits and upregulates Akt/mTOR signaling. **(A,B)** KEGG_ RIBOSOME pathway enrichment plots after 6 hours **(A)** and 72 hours **(B)** of actD vs. DMSO. NES, normalized enrichment score. **(C)** KEGG_RIBOSOME gene set enrichment plot for RNA expression after 72 hours of actD vs. DMSO. **(D,E)** KEGG_ PROTEASOME pathway enrichment plots after 6 hours **(D)** and 72 hours **(E)** of actD vs. DMSO. **(F)** KEGG_PROTEASOME gene set enrichment plot for RNA expression after 72 hours of actD vs. DMSO. **(G,H)** Phosphorylation of Akt/mTOR signaling components visualized and quantified by Western blot in WiT49 and 17.94 following 2nM actD. **(I)** Rapamycin kill curves for anaplastic Wilms tumor cell lines WiT49 and 17.94 (four-point regression line with shaded region representing 95% confidence interval). **(J)** Heat maps of Loewe synergy scores for combinations of actD and rapamycin in WiT49 and 17.94 cells.

**Supplementary Figure S5.**
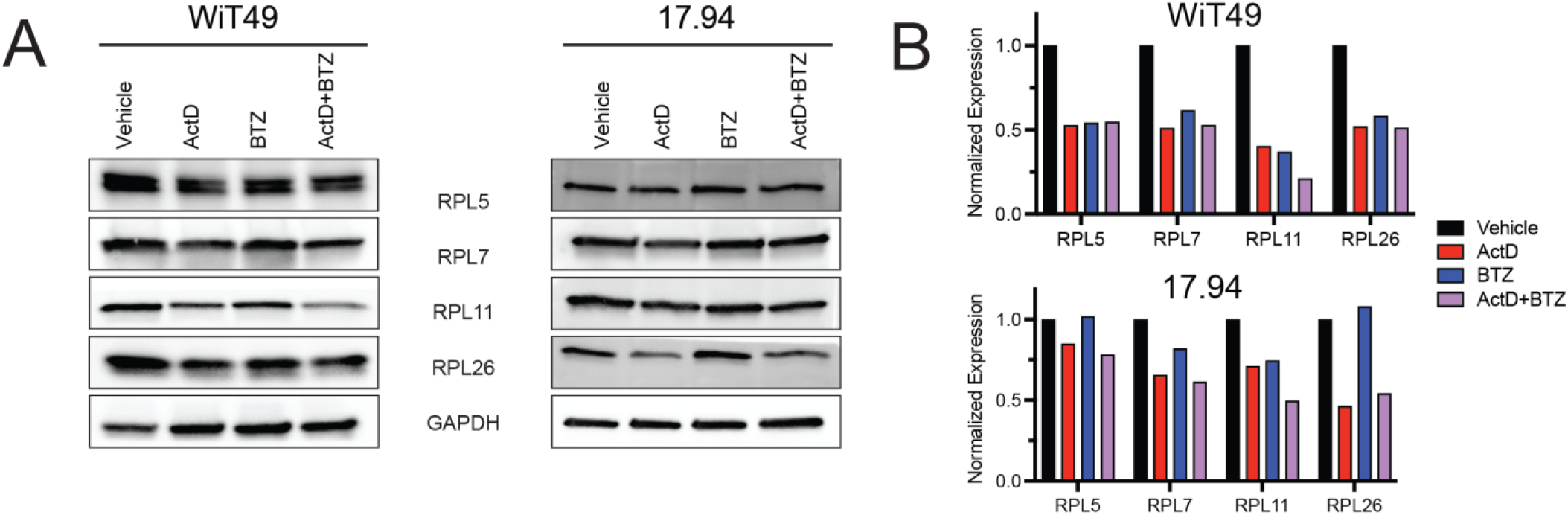
ActD and BTZ affect ribosomal protein levels. **(A)** Western blots for and quantification of **(B)** ribosomal protein subunits RPL5, RPL7, RPL11, and RPL26 in WiT49 and 17.94 following 48-hour treatment of DMSO, 2nM actD, 8nM BTZ, or 2nM actD + 8nM BTZ.

**Supplementary Figure S6.**
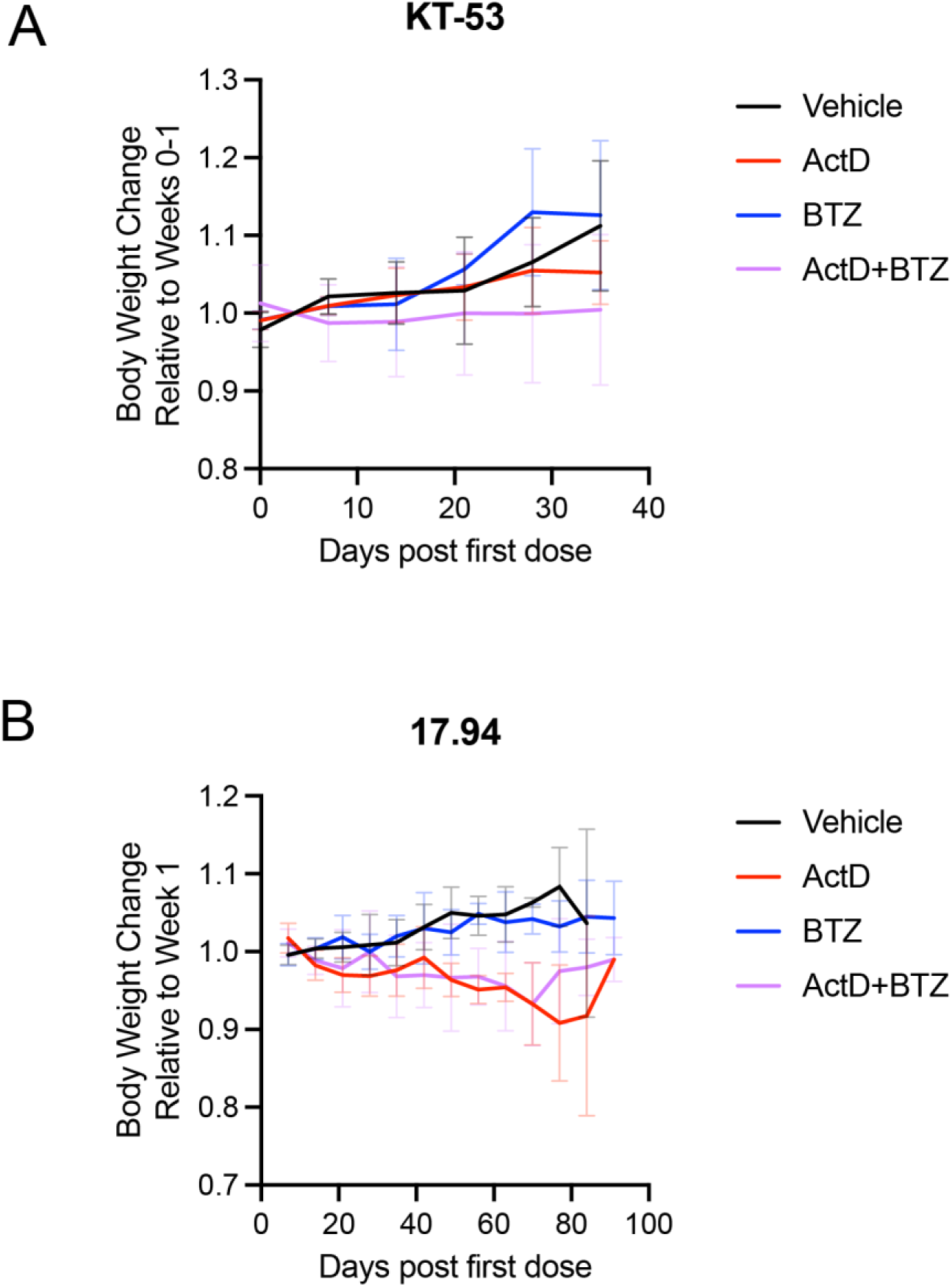
Body weight changes in mice treated with actD, BTZ, or combination. **(A,B)** Relative body weight of mice bearing KT-53 xenografts **(A)** or 17.94 xenografts **(B)**, during treatment with actD, BTZ, or both.

**Supplementary Figure S7.**
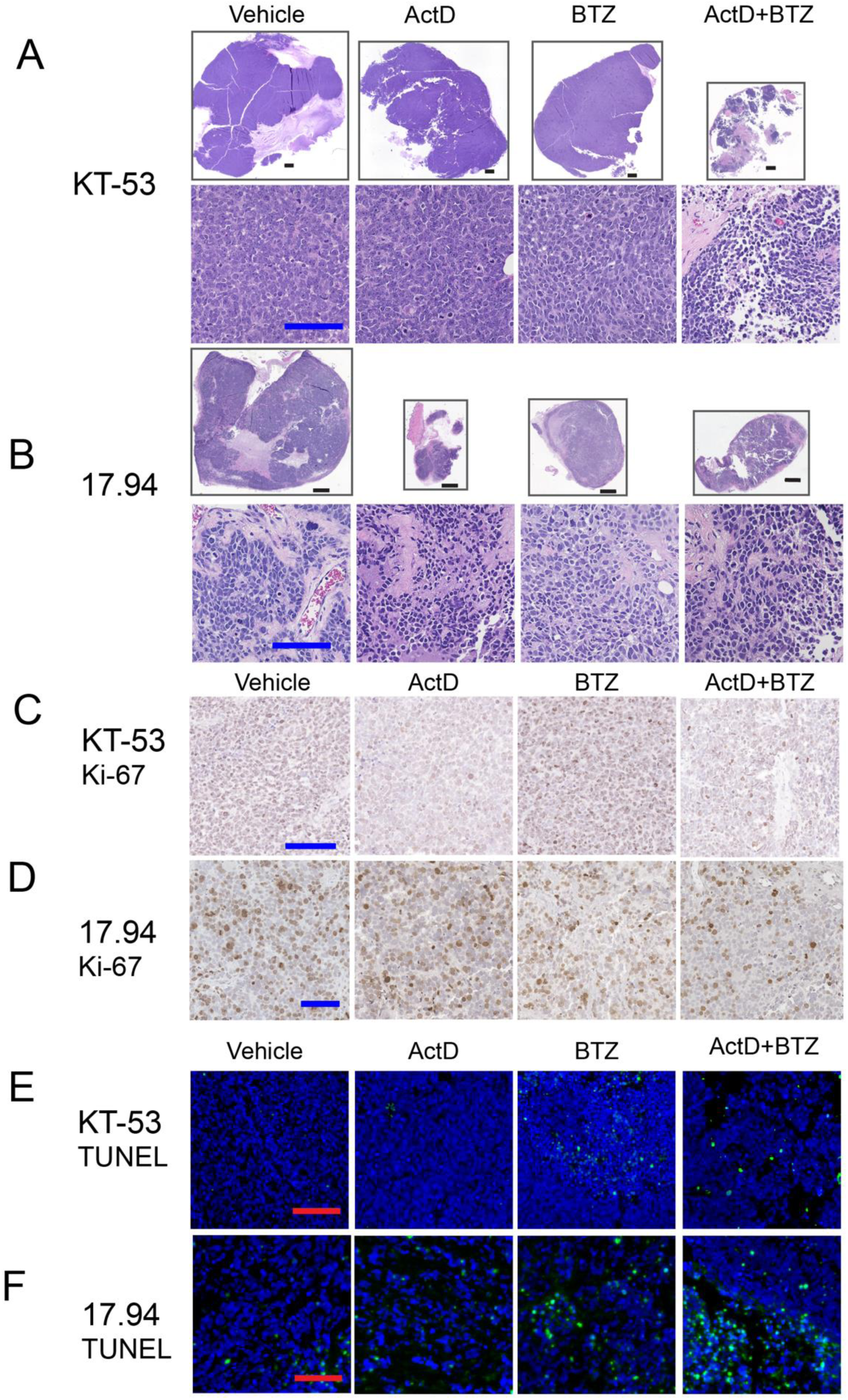
Bortezomib enhances cytotoxicity of actinomycin D in anaplastic Wilms tumor xenografts. **(A,B)** Low-power and high-power H&E images of **(A)** KT-53 and **(B)** 17.94 subcutaneous tumors (Black bar = 1.0 mm; Blue Bar =100 µm) **(C, D)** High-power photomicrographs of Ki-67 immunohistochemistry of KT-53 (Bar =100 µm) **(C)** and 17.94 **(D)**. **(E, F)** High-power photomicrographs of TUNEL stain of KT-53 (Bar =100 µm) **(E)** and 17.94 **(F)**.

**Supplementary Figure S8.**
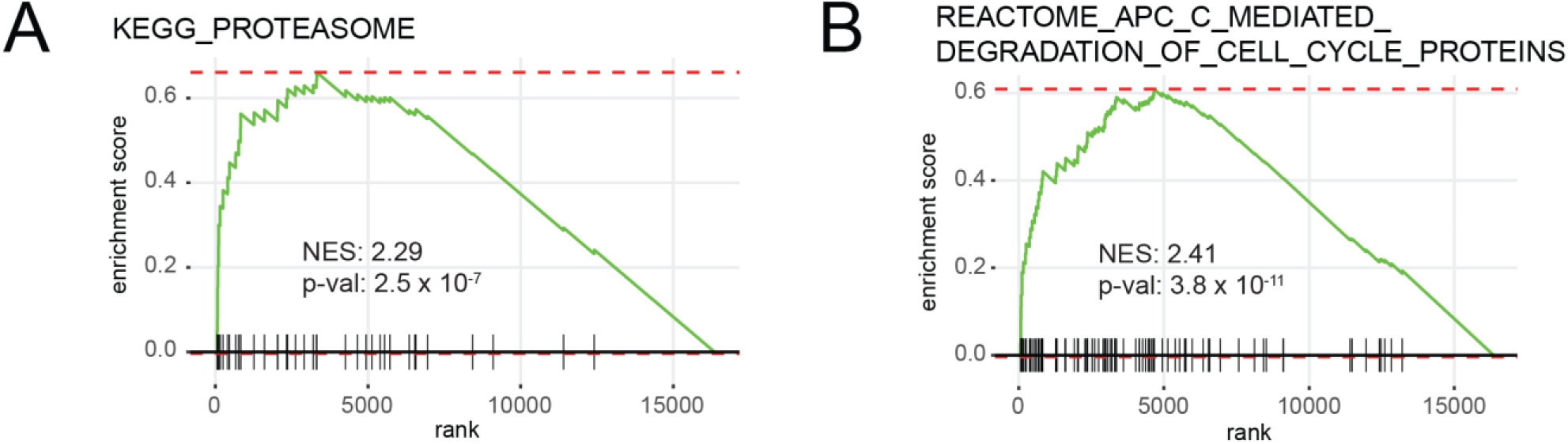
Enrichment for proteasome gene sets in anaplastic Wilms tumor. **(A,B)** GSEA plot showing enrichment for KEGG_PROTEASOME **(A)** and REACTOME_APC_C_MEDIATED_DEGRADATION_OF_CELL_CYCLE_PROTEINS **(B)** in anaplastic Wilms tumor vs. relapsed favorable-histology Wilms tumors in TARGET dataset.

**Supplementary Figure S9.**
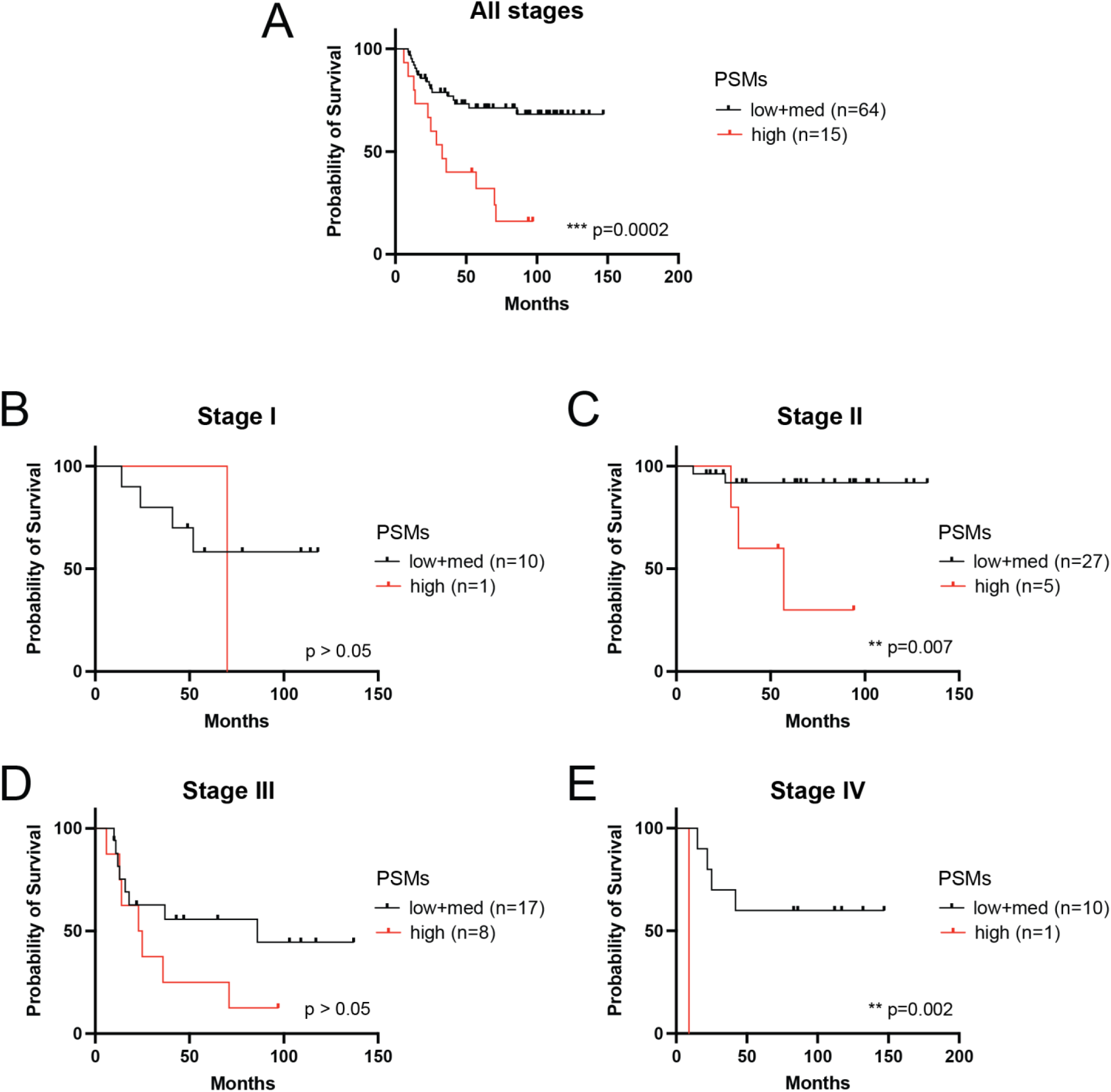
Survival of relapsed favorable histology Wilms tumor (FHWT) by stage and proteasome expression. **(A)** Overall survival of all relapsed FHWT in TARGET, stratified according to expression of proteasome enzymatic subunit genes *PSMB1*, *PSMB2*, *PSMB5*. **(B-E)** Overall survival of relapsed FHWT in TARGET, for Stages I, II, III, and IV at diagnosis, stratified according to expression of proteasome enzymatic subunit genes *PSMB1*, *PSMB2*, *PSMB5*. (Log rank test p-value: *<0.05, ** < 0.01, *** 0.001).

**Supplementary Figure S10.**
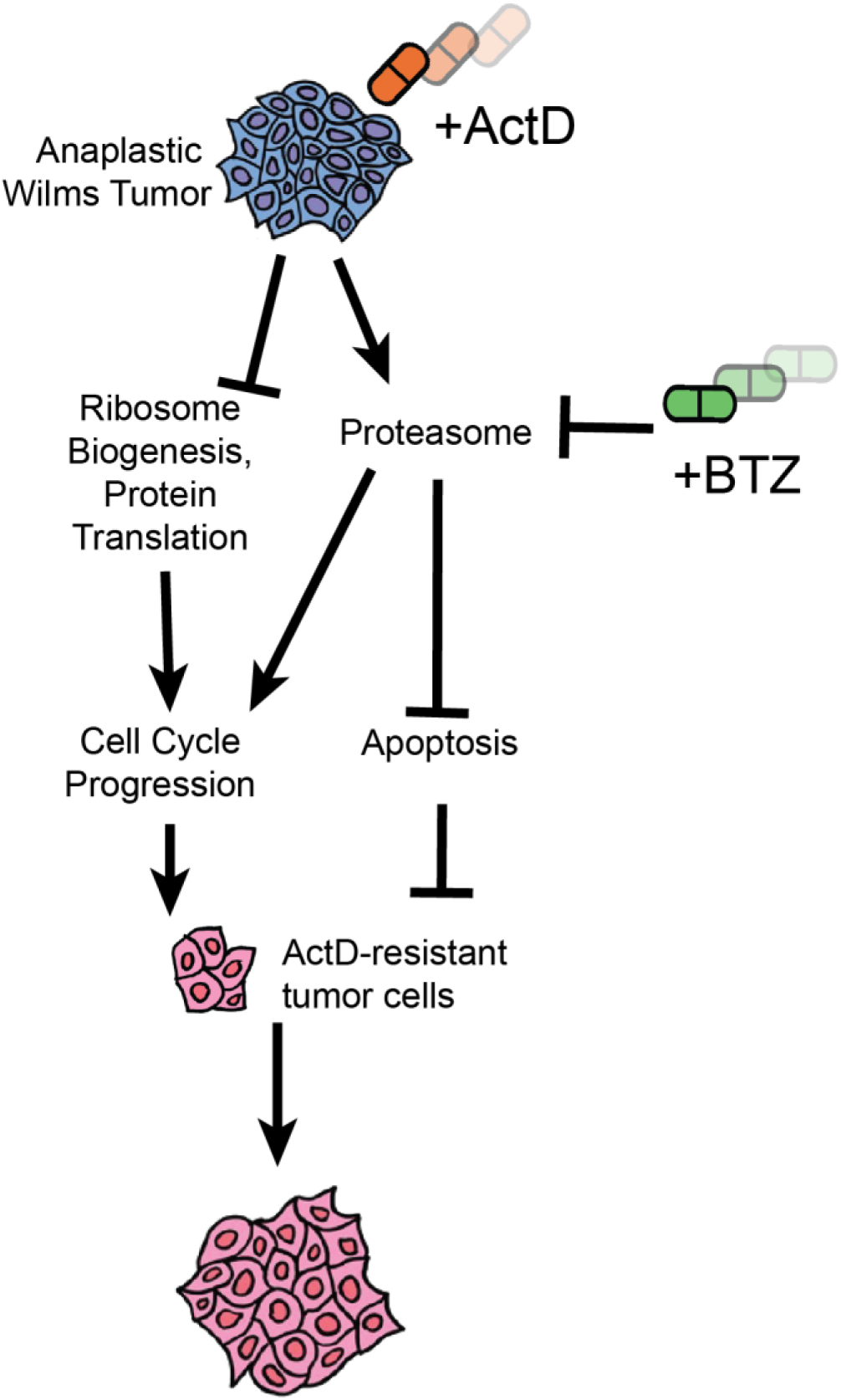
Model for effect of actinomycin D on protein homeostasis in Wilms tumor cells.

## SUPPLEMENTARY TABLES

**Supplementary Table S1.** Average log2-transformed translational efficiency for WiT49 in 6-hour and 72-hour actD versus DMSO treatments derived from ribosome profiling sequencing.

**Supplementary Table S2.** GSEA of translationally enriched KEGG pathways for WiT49 6-hour and 72-hour actD versus DMSO treatments from ribosome profiling sequencing.

**Supplementary Table S3.** KEGG pathway enrichment summary table from Mageck-Flute pathway output.

**Supplementary Table S4.** Normalized, log2-transformed, median-centered RPPA values of validated antibodies from WiT49 cells treated with actD or DMSO.

**Supplementary Table S5.** Loewe synergy scores (and 95% confidence interval) for actD with BTZ or rapamycin, in WiT49 or 17.94.

**Supplementary Table S6.** GSEA of enriched KEGG and Reactome pathways in DAWT versus FHWT.

**Supplementary Table S7.** List of oligonucleotides used in this study

